# Cellohood: multi-granular discovery of cellular neighborhoods with a permutation-invariant set transformer auto-encoder

**DOI:** 10.1101/2025.11.10.687184

**Authors:** Marcin Możejko, Daniel Schulz, Krzysztof Gogolewski, Nils Eling, Joanna Krawczyk, Olga Bogatyrova, Jakub Golab, Dominika Nowis, Robin Liechti, Marie Morfouace, Henoch Hong, Bernd Bodenmiler, Eike Staub, Ewa Szczurek

## Abstract

Discovering cellular neighborhoods and their roles in disease requires computational methods that consider the full breadth of data and offer multi-level granularity and interpretability. Here, we introduce Cellohood, a permutation-invariant set transformer auto-encoder equipped with a clinical association pipeline. Cellohood encodes full readouts for bags of spatially co-localized cells and supports multi-level analyses, providing interpretability by mapping latent dimensions to spatial tissue features. Our model surpasses current methods in accuracy of cellular neighborhood detection for spatial transcriptomics of the human cortex and CODEX spleen data measuring lupus progression. Applied to cancer data across three granularity levels, our method recovers neighborhoods along the expected immune-cold to immune-hot spectrum and further refines them into biologically and clinically meaningful subclasses, revealing spatial patterns linked to prognosis, histology, stage, and tumor mutational burden, and uncovering subgroups that transcend standard classifications. Overall, Cellohood enables in-depth analysis of complex tissues, revealing clinically informative spatial neighborhoods.

## Main

The recent explosion of spatial tissue profiling methods has revolutionized biology and medicine [**1**, **2**], enabling identification of *cellular neighborhoods*, i.e., coherent subregions with specific composition of cells [**3**]. Cellular neighborhood analyses offer insights into the basic principles of general morphological structures of healthy tissues [**4**] and how they change through disease progression [**5**], including the investigation of the interplay between tumors and their microenvironments, with its clinical outcomes [**6**], shedding light into causative pathobiological mechanisms [**7**].

Typically, cellular neighbourhood identification proceeds by finding representations of entire sets of co-localized cells or spots (which we refer to as *cell bags*) and then clustering them. The whole information about a given cell bag is contained in its *full cell bag representation,* that is a table containing molecular readouts for each cell in the bag (**Fig. 1**A). For spatial proteomics, readouts correspond to expression of protein markers per cell, while for spatial transcriptomics, they refer to genes per cell/spot. *Cell type representations* indicate the cell type of each cell in the bag. The full and cell type representations are technically challenging to work with, due to their **varying dimensionality**, as different cell bags may contain different numbers of cells, and due to **permutation equivariance**, as there is no canonical ordering of cells within a cell bag. Existing methods employ simple approaches to representing cell bags, falling into two categories. UTAG [**8**], STlearn [**9**], and CellCharter [**10**] work with *aggregated cell bag representation*, applying statistical reduction functions (typically mean, minimum, and maximum) across readouts from all cells within the cell bag. The approaches of Goltsev *et al*. [**5**], Jackson *et al*. [**6**], CytoCommunity [**11**], Spatial LDA [**12**], and Schürch *et al*. [**13**] use *histogram cell type representation* corresponding to cell type frequency within each cell bag and as such **are dependent on error-prone cell type assignment**. While both the aggregated and histogram representations circumvent technical challenges of full or cell type representations, offering fixed-dimensionality and permutation-invariance, they **lose information** of exact marker levels per each cell that may be biologically relevant. On top of that, while real applications may require either coarse-grained analysis to identify major morphological substructures or fine-grained investigation of complete spatial tissue heterogeneity, most existing approaches are **not equipped with the means to control the level of granularity**. Finally, the majority of the existing methods are **not easily interpretable**, i.e., one cannot trace back to the major axes of variation discovered by the model and interpret them visually to learn what the model took into account when defining the spatial neighborhoods.

**Figure 1.**
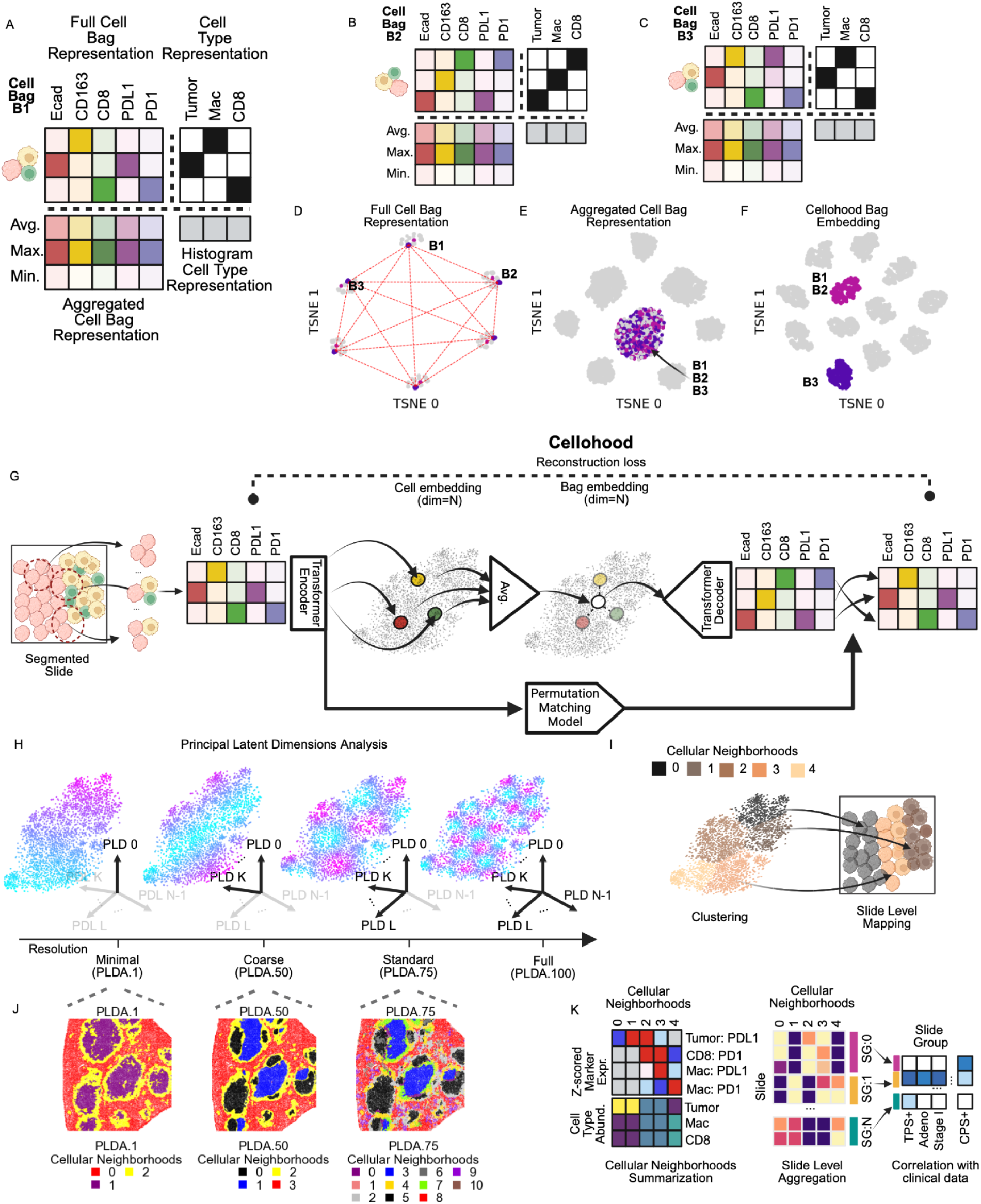
Conceptual overview of Cellohood. **(A)** Comparison of different cell bag representation methods on an example cell bag B1. Full cell bag representations store complete marker expression data. Cell type representations are reliant on cell type assignment. Both suffer from permutation dependence. Aggregated representations (Avg, Max, Min) lose individual cell information, while histogram representations only preserve cell type assignments. **(B)** Example cell bag B2 that differs from B1 only by the order of cells, yielding different full and cell type cell bag representation, but the same aggregated cell bag and cell type histogram representations. **(C)** Example cell bag B3, different from B1 and B2, with a macrophage that has an increased PDL1 expression instead of a tumor cell, but has the same aggregated cell bag and histogram cell type representations. **(D)** T-SNE visualization of simulated cell bags, with different permutations of the same bag represented as different objects, demonstrating the permutation dependence problem. **(E) T**-SNE visualization demonstrating the information loss of aggregated cell bag representation. **(F)** Cellohood bag embeddings maintain both permutation invariance and cell-specific information, properly clustering similar bags (B1, B2) while distinguishing different ones (B3). **(G)** Cellohood architecture showing the permutation-invariant set transformer auto-encoder framework with permutation matching model. The model creates cell embeddings and bag embeddings while maintaining reconstruction ability of the original marker expression matrices. (**H**) Principal Latent Dimensions Analysis (PLDA) at different resolutions. PLDA.1 represents a single PCA dimension, PLDA.50 uses PCA dimensions explaining 50% of variance, PLDA.75 - 75% of variance, and PLDA.100 **-** all dimensions, showing increasing resolution from minimal to full. **(I)** Segmented slides at preselected PLDA resolution are processed into cell bags, which are embedded by Cellohood and clustered into cellular neighborhoods. (**J**) Visualization of different cellular neighborhoods slide level mappings for an example slide from CODEX Spleen dataset from [**5**] showing increasing number of identified neighborhood types with higher resolution settings, revealing finer structural patterns in the tissue. (**K**) Clinical association pipeline. After summarization w.r.t. to cell type abundance and marker expression profiles, cellular neighborhoods are aggregated at a slide level for correlation with clinical data.

Here, to close those important gaps, we propose Cellohood, a permutation invariant set transformer auto-encoder model for the discovery of cellular neighborhoods. Cellohood models entire cell bags of varying sizes in a way that preserves the full readout information, without reliance on cell type assignment. The model offers the opportunity to operate on different levels of granularity of data representation, thereby allowing for the inference of major morphological substructure (coarse grained level) as well as detailed microenvironments (full detail level). We demonstrate that the latent dimensions of the model map back to spatial features of the analysed data, offering high interpretability. Finally, we equip the Cellohood framework with a clinical association analysis tool, which enables exploration of identified spatial neighborhoods and statistical testing of associations with clinical features. We demonstrate the capabilities of Cellohood by benchmarking the model on two datasets with ground truth: the spatial transcriptomics dataset for human prefrontal cortex [**4**] and CODEX dataset profiling spleen from lupus mice [**5**]. To showcase Cellohood as a novel discovery tool with different granularity levels, we apply it to 1307 IMC slides across 2 marker panels for 191 non-small cell lung cancer (NSCLC) patients that we generated as part of the IMMUcan consortium [**14**, **15**], as well as a previously published breast cancer cohort [**6**].

## Results

### Cellohood overcomes limitations of cell bag representations

To empirically demonstrate the technical challenges related to the full cell bag representation and the limitations of the alternative representations, we considered three example cell bags (Fig. 1A-C). Different cell ordering yielded different full cell bag representations (compare Fig. 1A to Fig. 1B). On the other hand, identical cell type representations were obtained for cell bags differing in biologically relevant marker PDL1 (compare Fig. 1A to Fig. 1C). Similarly, the histogram cell type representations were identical across all cell bags due to the fixed cellular composition, entirely masking PDL1 expression patterns (Fig. 1A–1C). The aggregated representation approach also failed to distinguish between biologically different immune microenvironments (Fig. 1A–1C). To further illustrate issues with information loss, we generated a simulated proteomics dataset of cell bags, each containing exactly three cells expressing five markers: Ecad, CD163, CD8, PD1, and PDL1 (Extended Fig. 1, Extended Text Section 1). For this dataset, biologically identical simulated cell bags with different cell orderings clustered separately (Fig. 1D). On the other hand, clustering of aggregated representations of different cell bags grouped them in shared clusters (Fig. 1E). In contrast, the cell bag embeddings obtained by the proposed Cellohood approach preserve the full biological information contained in the full cell bag representations, while gaining the advantage of permutation invariance and fixed dimensionality characteristic of the aggregated or histogram representations (Fig. 1F).

### Cellohood Overview

Cellohood is a permutation-invariant set transformer auto-encoder model that aims at providing new bag-informed cell embeddings and entire bag embeddings that preserve information about the features of each individual cell (see Fig. 1G). After segmentation, identified cells in a given slide are spatially clustered into cell bags. The clustering is based on physical proximity, and depending on tissue density yields different numbers of cells per bag. The Cellohood encoder accepts as its input a full cell bag representation and returns a new, fixed-size cell embedding for each cell in the bag. The permutation invariance of each of these embeddings is guaranteed by the permutation invariance of a vanilla transformer. These embeddings are then averaged to provide a bag embedding, which is also guaranteed to be fixed size and permutation invariant, as the average function is permutation invariant. Subsequently, the bag embedding is fed to a decoder in order to reconstruct the original cell bag representation. Because of the permutation invariance, the permutation of the decoder output might differ from that of the input cell bag. To alleviate this problem, an additional Permutation Matching model is trained jointly with Cellohood in order to match the permutation of the decoder output and the input bag.

To systematically control the resolution of spatial analysis within our framework, we developed Principal Latent Dimensions Analysis (PLDA), an approach for modulating the level of detail retained by the analyzed embeddings (see Fig. 1H). PLDA applies principal component analysis to cell or bag embeddings and selectively utilizes an implicitly defined number of principal components for cellular neighborhood (CN) determination. The number of components ranges from a minimal representation using only the first principal component (PLDA.1), to increasingly comprehensive representations that capture 50% (PLDA.50), 75% (PLDA.75), or 100% (PLDA.100) of the latent variation.

Following training, we leverage Cellohood for systematic identification of CNs within the analyzed tissue (Fig. 1I). The Cellohood cell and cell bag embeddings at a preselected PLDA resolution are subjected to unsupervised clustering, yielding distinct CNs that represent prototypical spatial configurations within the tissue architecture. The identified neighborhoods are then spatially mapped onto the original tissue sections, enabling direct comparison with established histological features and tissue compartments.

Fig. 1J demonstrates the biological relevance of PLDA multi-resolution approach using CODEX imaging data from mouse spleen [**5**]. As resolution increases from PLDA.1 to PLDA.75, slide level mapping shows progressively finer neighborhood structures, revealing distinct organizational patterns that would remain obscured at lower resolutions. The minimal resolution (PLDA.1) identifies only fundamental tissue compartments (white pulp, red pulp and a marginal zone between them), while higher resolutions (PLDA.50, PLDA.75) reveal intricate cellular communities with distinct marker profiles in increasing detail. This scalable approach allows researchers to select the appropriate resolution for their specific biological question, balancing between coarse to fine-grained tissue patterns.

To enable discovery of associations between CN structure and clinical features, we developed a CN clinical association pipeline for quantitative characterization of identified CNs by their cell type abundance and expression profiles, whole-slide level clustering by aggregated CN abundances, revealing distinct slide groups (SGs) suitable for quantifying association with sample-level metrics and clinical parameters using Benjamini–Hochberg corrected one-sided hypergeometric test (Fig. 1K).

### Cellohood outperforms other models in identifying human dorsolateral prefrontal cortex layers and its latent embeddings organize according to the layers

To demonstrate the efficacy of Cellohood on a benchmark dataset with expert-defined spatial neighborhoods, we performed an analysis of the spatial transcriptomics Visium 10X human dorsolateral prefrontal cortex (DLPFC) dataset, using the PLDA.50 coarse resolution cell embedding. This resolution was chosen to enable capturing major morphological structures, which we expected to correspond to the DLPFC layers. With the Adjusted Rand Index (ARI) of 0.58, Cellohood showed superior performance in retrieving the expert-defined DLPFC layers (Fig. 2A) as compared to state-of-the-art methods, including Cellcharter [**10**], STAGATE [**16**], BayesSpace [**17**], SEDR [**18**], DR [**19**], UTAG [**8**], and the baseline approach of clustering aggregated cell bag representations, referred to as Agg.CBR [**20**] Interestingly, the simple Agg.CBR emerged as the third most effective approach, surpassed only by Cellohood and Cellcharter. We did not include a comparison to a clustering of histogram representations, as cell types for this dataset were defined based on the layer information.

**Figure 2.**
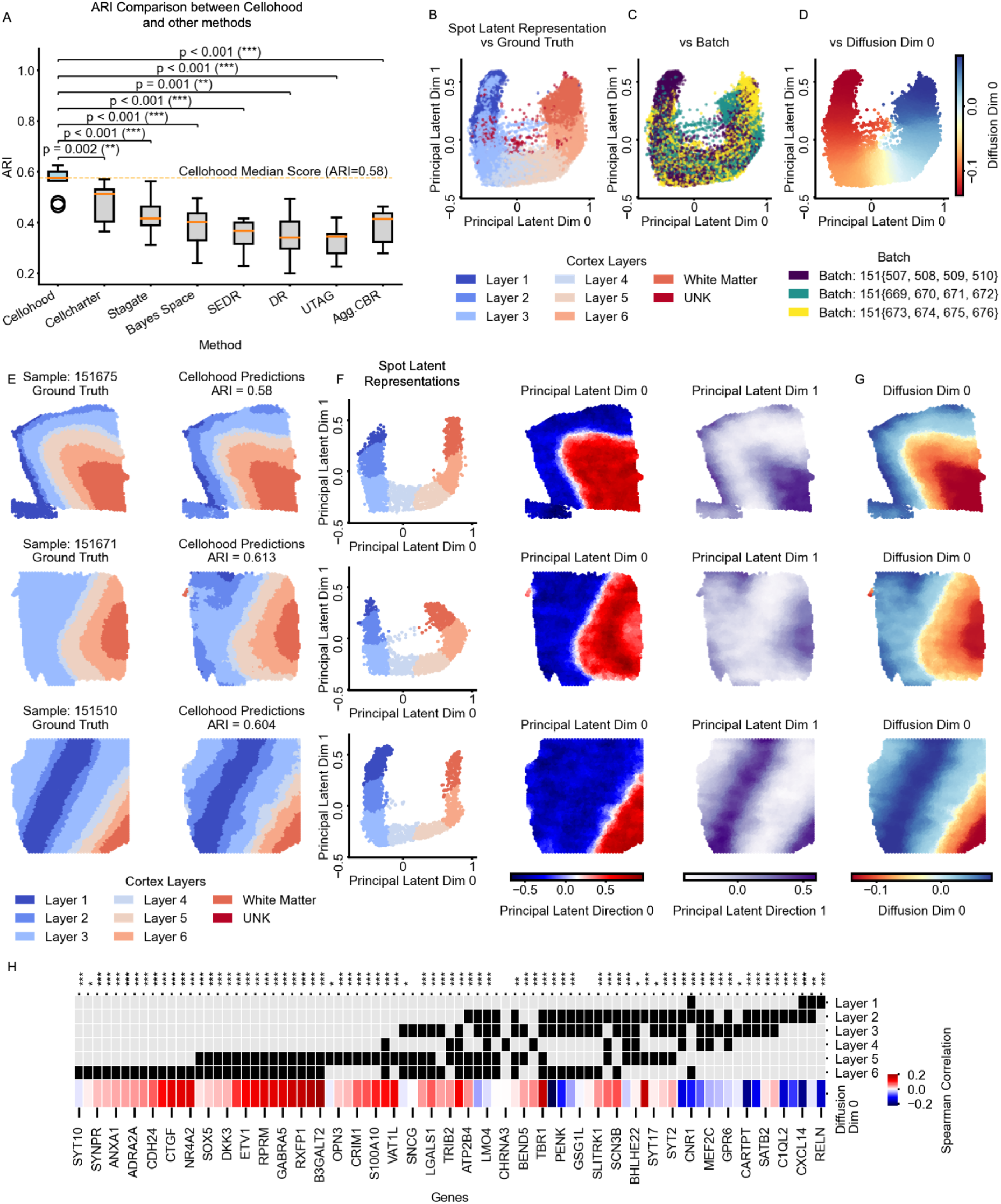
Performance of Cellohood on Visium10X DLPFC dataset. (**A**) Quantitative comparison of classification accuracy using Adjusted Rand Index (ARI) across multiple methods. Statistical significance of performance differences measured by two-sided paired t-Test is indicated by asterisks (*p≤0.05, **p≤0.01, ***p≤0.001). (**B**-**C**) PLDA.50 representations of tissue spots revealing clear separation of cortical layers (color-coded) (**B**) and batch effects (**C**). (**D**) Diffusion Dimensionality Reduction on PLDA.50. (**E**) Representative samples (151675, 151671, 151510; rows) from each patient in the study, with ground truth annotations and Cellohood predictions. (**F**-**G**) PLDA.50 based representations provide insight into the model’s internal feature extraction. (**H**) Spearman correlation (colors) between Diffusion Dim 0 and gene expression of marker genes of cortical layers from [**22**] and [**4**], with marker gene status for each layer encoded as black entries.

Further investigation of the Cellohood latent embeddings revealed a distinctive U-shaped topology in the DLPFC dataset that corresponded to the cortical layer organization (Fig. 2B). This structure persisted across batches, indicating robustness of our model to technical variation (Fig. 2C). To better characterize this topological feature of Cellohood latent embeddings, we employed Diffusion Dimensionality Reduction [**21**], which effectively captured the U-shaped pattern in its first dimension (Diffusion Dim 0; Fig. 2D). This analysis illustrates that the Cellohood latent embedding space effectively captures the underlying biological continuum of cortical layers.

Detailed investigation of three representative samples (151675, 151671, and 151510), each from a distinct patient, revealed high agreement of Cellohood predictions with ground truth, mapping into distinct spot neighborhoods within the spatial regions marked by the cortex layers (Fig. 2E; see Extended Fig. 2A-F for all samples). The spot latent representations are arranged in neighborhoods in agreement with the consecutive layers and are reflected in the values of Principal Latent Dimensions 0 and 1 (Fig. 2F), as well as Diffusion Dimension 0 (Fig. 2G), confirming that the model captures genuine biological structure rather than technical artifacts.

Spearman’s correlation analysis between Diffusion Dimension 0 and established marker genes [**4**, **22**] further validated the biological significance of the identified U-shaped pattern (Fig. 2H, Extended Fig. 2G). The significant correlations observed between this dimension and canonical layer-specific transcriptional signatures provide molecular evidence that Cellohood effectively captures the underlying biological organization of cortical architecture.

### Cellohood recapitulates known and introduces finer detail in neighborhood analysis of lupus-affected spleen structure

We next evaluated Cellohood against leading computational methods, including CytoCommunity [**11**], SpatialLDA [**12**], UTAG [**8**], STAGATE [**16**], BayesSpace [**17**], and stLearn [**9**], as well as two baselines, Agg.CBR (clustering aggregated cell bag representations) and CTH.CBR (clustering histogram cell type representations) [**20**] in application to spatial proteomics CODEX data from healthy and lupus-affected murine spleen samples [**5**]. In this analysis, we again used PLDA.50 coarse resolution, which we expected to indicate major spatial patterns characteristic of the disease progression. Using Cellohood representations of the healthy samples achieved mean scores of F1=0.76 and Adjusted Mutual Information (AMI)=0.58, demonstrating superior accuracy in identifying expert-defined architecture of splenic compartments (B-zone, periarterial lymphatic sheaths (PALS), marginal zone, and red pulp; Fig. 3A).

**Figure 3.**
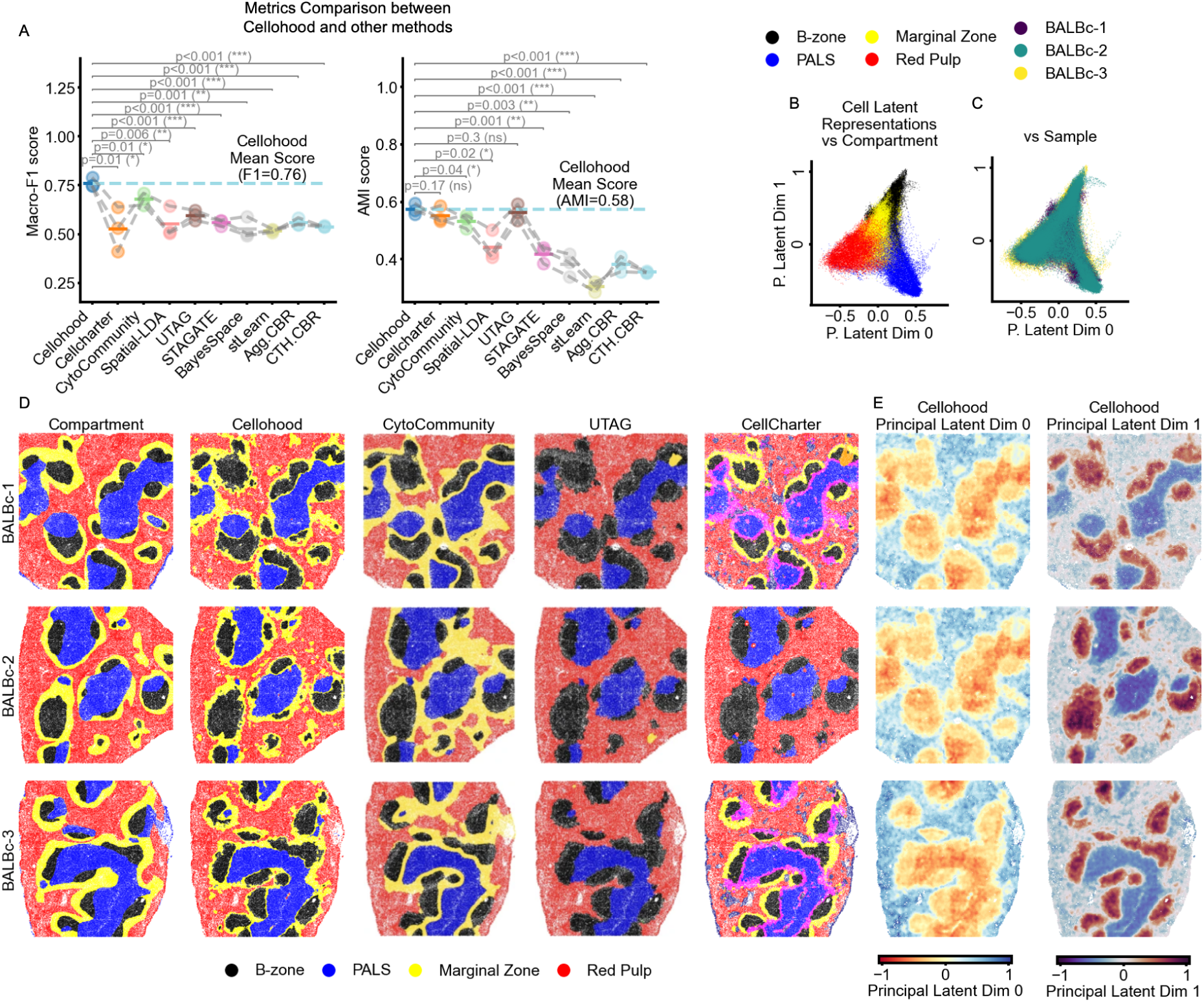
Cellohood performance on CODEX Spleen data [5]. **(A)** Comparison of Cellohood **to** other methods using F1 score (left) and Adjusted Rand Index (ARI) (right). Statistical significance measured by one-sided t-Test, indicated by asterisks (*p≤0.05, **p≤0.01, ***p≤0.001)**. (B-C)** PLDA.50 representations of cells showing clear separation of splenic regions (B-zone, PALS, Marginal Zone, Red Pulp) (**B**), and lack of clear differences between 3 healthy samples (BALBc-1-BALBc-3) (**C**). **(D)** Ground truth annotations (left column) compared with predictions from Cellohood (second column), CytoCommunity (third), and UTAG (fourth) across three BALB-c samples (rows). **(E)** Cell latent representations and first two principal latent dimensions from Cellohood.

Analysis of the first two Principal Latent Dimensions (0 and 1) of Cellohood revealed distinct clustering of cell neighborhoods corresponding to key splenic compartments, separating B-zone, PALS, marginal zone, and red pulp regions (Fig. 3B, Extended Fig. 2A). Notably, these Latent Dimensions were not impaired by batch effects from sample of origin (Fig. 3C, Extended Fig. 3B) indicating that Cellohood effectively captures fundamental tissue architecture independent of technical variation across samples.

We further applied slide-level mapping of predicted compartments to provide visual validation of Cellohood’s performance through side-by-side comparisons of ground truth annotations with computational predictions across three healthy mouse spleen samples (Fig. 3D). Cellohood’s predictions closely match the expert-annotated ground truth, accurately identifying boundaries between distinct splenic regions, while both baseline methods (Agg.CBR and CTH.CBR) [**20**] failed at this task (Extended Fig. 3C,D). Interestingly, UTAG [**8**] completely missed to identify the marginal zone despite achieving AMI scores comparable to Cellohood, highlighting the limitation of relying solely on quantitative metrics for method evaluation. Similarly, Cellcharter failed to predict the marginal zone in one sample (BALBc-2) while introducing spurious PALS-B-zone marginal zones and sub-compartments of red pulp, yet also achieved comparable AMI scores. Finally, CytoCommunity [**11**] tends to overpredict the marginal zone, resulting in less accurate boundary determination. Cellohood’s cell latent representations and principal latent dimensions mapped to tissue coordinates for the same samples indicate that Cellohood not only provides accurate domain labels but also learns biologically meaningful representations that reflect the underlying spatial structure of the spleen, particularly for challenging regions like the marginal zone that other methods struggle to identify accurately (Fig. 3D).

To explore additional insights from higher resolution analysis and full dataset, we applied Cellohood PLDA.100 on the full set of samples (including healthy and lupus affected), finding that lupus progression is marked by loss of normal splenic compartmentalization, expansion of activated T- and CD5⁺ B-cell neighborhoods, disruption of the marginal zone, and emergence of chaotic inflammatory niches (Extended Fig. 3F-I, Extended Text Section 2).

### Minimal resolution analysis of the IMMUcan NSCLC dataset reveals major axes of spatial variation in IMC data

We next deployed Cellohood to discover spatial tumor microenvironmental patterns and their associations with clinical characteristics in lung cancer patients. Through the IMMUcan consortium, we generated IMC data using two marker panels (Panel 1: 40 immune profiling markers (Extended Table 1); Panel 2: 44 tumor microenvironment markers (Extended Table 2) and collected clinical data for 192 NSCLC patients (Extended Tables 3,4). This versatile dataset enabled us to showcase the types of insights obtainable about spatial patterns of tumors from non-coarse PLDA resolutions: minimal, standard and full, which were not used in the analyses of benchmark datasets.

We first employed the minimal PLDA.1 to investigate Panel 1 data, with up to four IMC slides per patient: three from distinct tumor areas and one from CD20+ areas (tertiary lymphoid structure (TLS) proxy) for 172 patients, totaling 741 slides. Cellohood was trained on cell bags from all slides.

For tumor slides, PLDA.1 cell embedding analysis revealed six distinct cellular neighborhoods (CNs) representing increasing immune infiltration levels (Fig. 4A, Extended Fig. 3A, Extended Table 5). CN 0 comprised immune-cold, tumor-only regions, while CN 6 represented immune-hot neighborhoods. Ki67 proliferation marker expression was highest in immune-cold CN 0 and progressively decreased in other CNs, while active immune response markers peaked in CN 6 (Fig. 4B). CN 2 was characterized by mural cells showing highest SMA expression, indicating activated cancer-associated fibroblasts (CAFs). Correlation analysis confirmed that CNs with similar immune infiltration profiles were highly correlated (Fig. 4C), supporting the biological relevance of Cellohood CNs.

**Figure 4.**
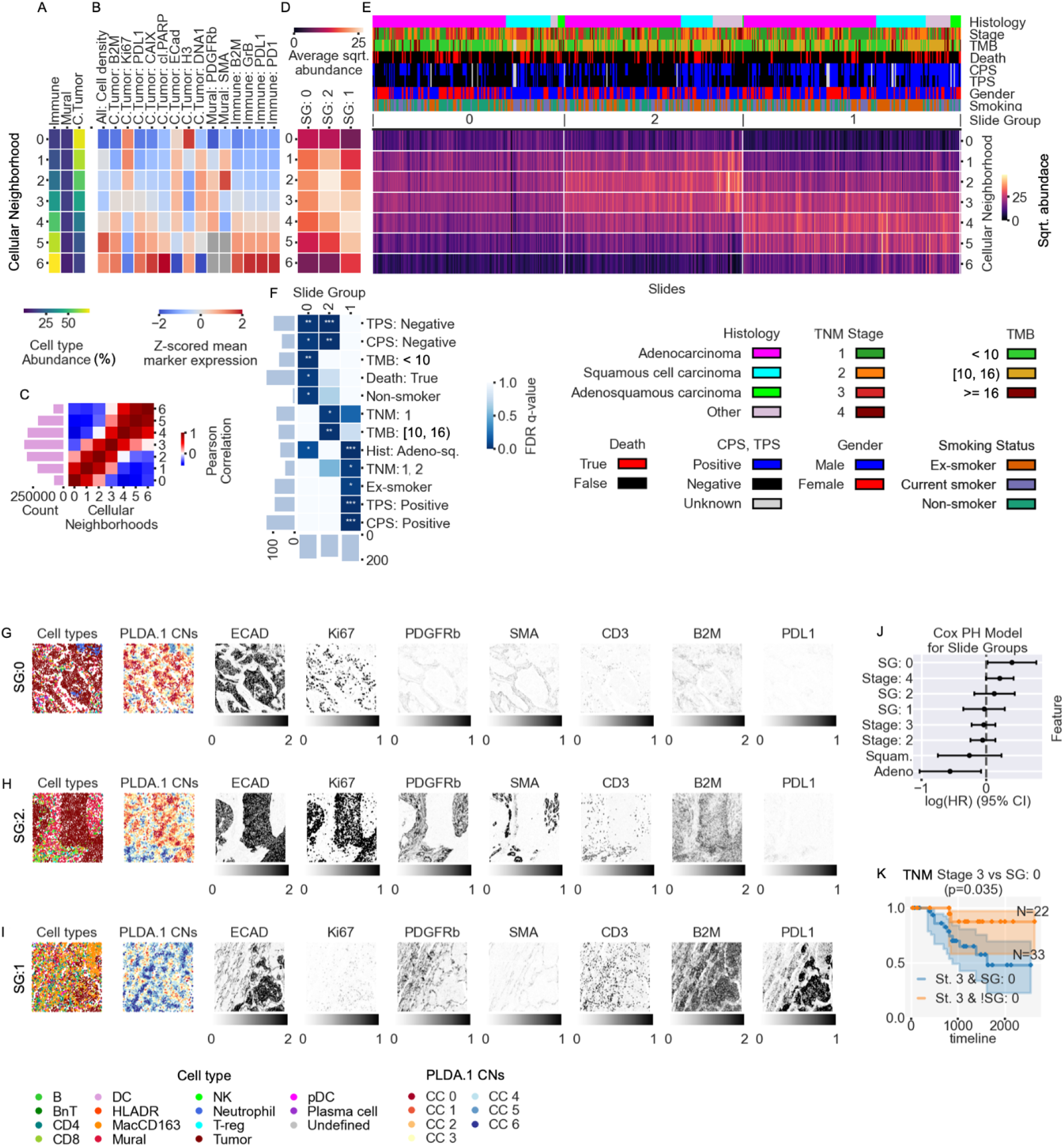
Minimal resolution (PLDA.1) cell embedding-based cellular neighborhoods (CNs) and their clinical relevance for IMMUcan NSCLC IMC Panel 1 dataset. (**A**) Average cell class abundances (columns) for CNs (rows). (**B**) Z-scored average marker expressions across specific or all cell classes (columns) for CNs (rows), presented only for cell classes comprising >10% of a given cluster. (**C**) Correlation between CN abundances at the slide level. Side bars: total CNs count across the cohort. (**D**) Slide groups (SGs; columns), characterized by average square rooted abundance of CNs (rows). (**E**) Abundances of CNs (rows) for individual slides (columns). Colors in top rows: clinical covariates associated with each patient. (**F**) Association of SGs (columns) with clinical covariates (columns), with FDR-corrected q-values from one-sided hypergeometric tests performed for each SG against each clinical variable (∗q≤0.2, ∗∗q ≤0.05, ∗∗∗q≤0.01). (**G**-**I**) Representative slides from SG 0 (G), SG 1 (**H**) and SG 2 (**I**), with cell types and selected marker expressions. (**J**) Multivariate Cox proportional hazard model for SGs and relevant clinical covariates, and survival of TNM Stage 1 patients established as the baseline. Error bars represent 95% confidence intervals for each hazard ratio. (**K**) Kaplan-Meier curves for TNM Stage 3 patients stratified by SG 0.

Following the clinical association pipeline (Fig. 1K), we aggregated the identified PLDA.1 CN abundances at slide level and clustered them into three slide groups (SGs) (Fig. 4D,E), which we analyzed for clinical associations (Fig. 4F). Both SG 0 (with a representative slide presented in Fig. 4G, which featured an immune-cold landscape, dominated by CNs with high tumor and low immune cell type abundance) and SG 2 (representative in Fig. 4H, with high abundance of tumor-immune border CNs 2 and 3), were significantly associated (hyper geometric test q-value < 0.05) with low Tumor Proportion Score (TPS; indicative of high PDL1 expression) [**23**]. At the same time, SG 0 was strongly enriched with low tumor mutational burden (TMB) [**24**], while SG 2 with high TMB patients. SG 1 (representative in Fig. 4I, capturing immune-infiltrated regions) was strongly enriched in patients with high CPS/TPS and rare adeno-squamous histology. Cox Proportional Hazard modeling revealed that patients with slides in immune-cold SG 0 had the highest hazard ratio (Fig. 4J). Moreover, SG 0 presence stratified Stage 3 patients into distinct prognostic groups with significantly different survival profiles (log rank test p=0.035; Fig. 4K). Thus, spatial patterns characteristic for SG 0 provide clinically relevant insights that extend beyond traditional tumor prognosis approaches. Taken together, this analysis validated established knowledge that lung tumors vary along a biological spectrum from immune-cold to immune-hot states, with immune-excluded regions showing high proliferation and poor outcomes.

### Standard resolution of the NSCLC dataset exposes finer grained cellular neighbourhoods

Next, to obtain more fine-grained neighborhoods, we analyzed all slides from this cohort using standard resolution (PLDA.75) cell embeddings, revealing a finer grained collection of 23 distinct CNs, which we subcategorized into 7 classes: *pure tumor* (tumor-dominated neighborhoods), *infiltrated* (consisting of a mix of tumor and immune cells), *CD20*+ (with a high abundance of B and B and T (BnT) cells), as well as *neutrophil*-, *mural*-, *macrophage*- and *plasma cell-enriched* neighborhoods (Fig. 5A,B). Among pure tumor neighborhoods, tumor cells showed distinct marker profiles: CN 13 (low marker expression), CNs 6 and 22 (high Ki67), CN 17 (increased PD-L1), CN 4 (high TCF7/E-cadherin with MHC class I), CN 10 (high hypoxia marker CAIX with MHC class I), and CN 18 (increased CD15). Neutrophil-enriched CN 8 showed tumor cells with elevated PD-L1 and apoptosis markers. Infiltrated neighborhoods featured tumor cells with high MHC class I (CN 21) or apoptosis markers (CN 9). CN 20 displayed co-stimulation (ICOS) and checkpoint (LAG3) molecules with prolonged antigen stimulation markers (CD45RO, PD-1) on T cells. Macrophage-enriched CN 12 showed macrophages with elevated CD68 and CD206 expression. Plasma cell-enriched neighborhoods CN 5 and CN 15 exhibited high VISTA expression across all cells, with CN 5 additionally showing LAG3 on all cells and CD45RA on T cells. In summary, this higher resolution analysis provided more detailed characterization than PLDA.1, capturing specific tumor cell phenotypes ranging from proliferative to hypoxic states, as well as specialized immune niches with unique activation and exhaustion profiles.

**Figure 5.**
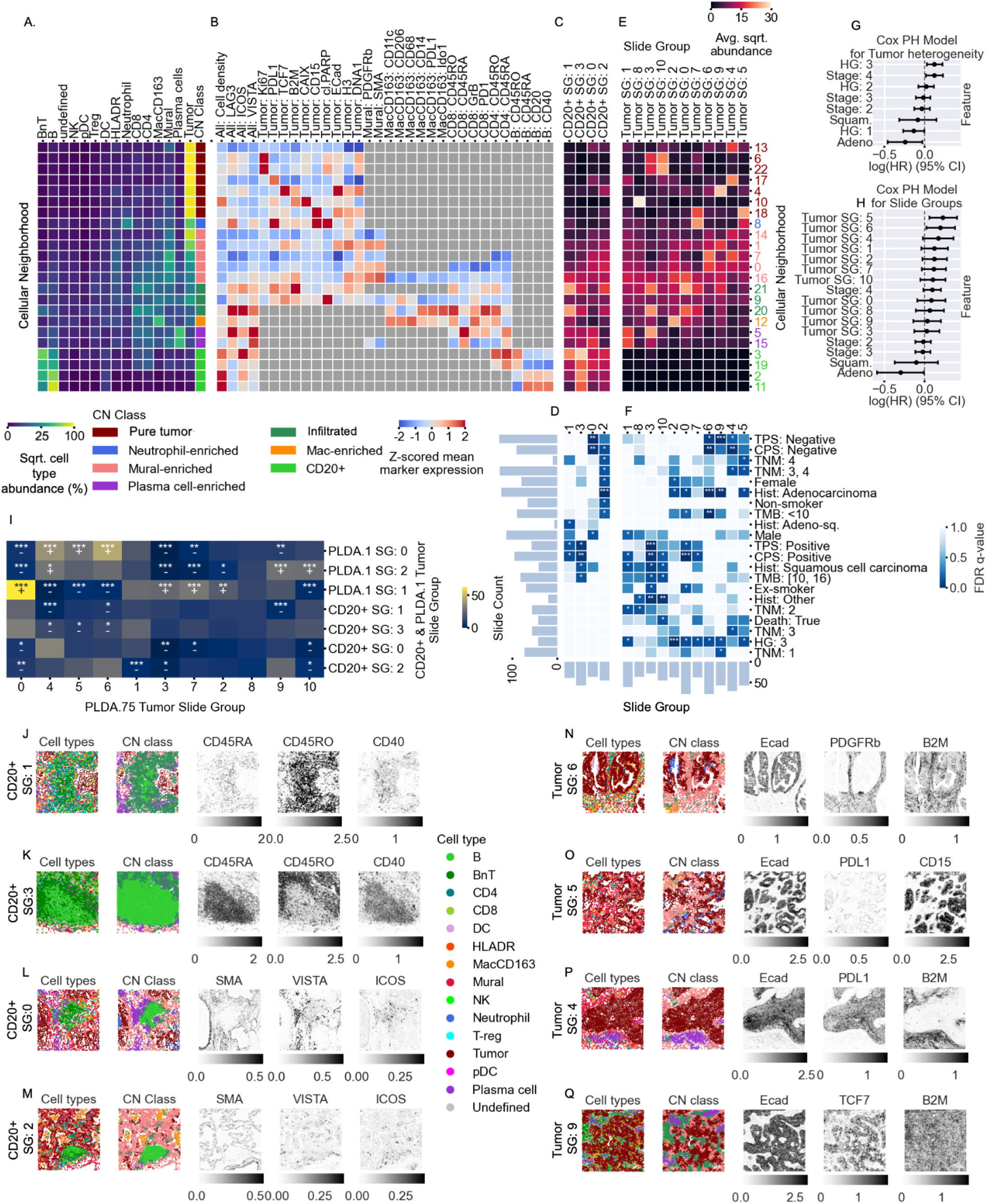
Cellohood Results on IMMUcan NSCLC Panel 1 dataset for PLDA.75 analysis. (**A**) Average cell type (rows) percentage for each CN (columns). (**B**) Z-scored average marker expressions across specific or all cell types (rows) for CNs (columns), presented only for cell types comprising >10% of a given cluster. (**C**-**D**) Slide groups (SGs; columns), characterized by square rooted abundance of CNs (rows) for CD20^+^ slides (**C**) and their association with clinical covariates (**D**). (**E**-**F**) Analogous slide groups (**E**) and clinical associations (**F**) for tumor slides. (**G**-**H**) Cox proportional hazard model for tumor heterogeneity classes (**G**) and tumor slide groups (**H**) with relevant clinical covariates, and survival of TNM Stage 1 patients established as the baseline. Error bars represent 95% confidence intervals for each hazard ratio. (I) Cross-counts of PLDA.75 Tumor slide groups with PLDA.1 and CD20^+^ PLDA.75 slide groups together with FDR-corrected q-values from two-sided hypergeometric tests (∗q≤0.2, ∗∗q≤0.05, ∗∗∗q≤0.01, + indicating enrichment, - indicating avoidance). (**J**-**N**) Representative slides from PLDA.75 CD20^+^ (**J**-**M**) and Tumor (**N**-**Q**) slide groups, with cell types and CN class and selected marker expressions.

In order to analyze CD20+ slides in PLDA.75 resolution, we clustered them into 4 groups based on slide-aggregated abundances and correlated them with clinical covariates (Fig. 5C,D, Extended Fig. 5A,B, Extended Table 6). SG 1 was characterized by high abundance of CNs 21 and 20 (representative slide in Fig. 5J). SG 3 showed TLS composition with crowded aggregations of B- and T-cells and high CD40 and CD45RA expression on B-cells (Fig. 5K), and was significantly associated with increased CPS. In contrast, SG 0 and SG 2 contained slides with fewer lymphocytes, predominantly in stromal areas (Fig. 5L,M). SG 0 showed higher abundance of plasma cell-enriched CNs 5 and 15, while SG 2 had more immune cell-infiltrated CNs 20 and 21. SG 0 was strongly enriched in patients with negative TPS and CPS, while the most significant association of SG 2 was with adenocarcinoma patients. Together, these findings highlight that TLS-associated neighbourhoods rich in B–T cell interactions (SG 3) align with increased, whereas plasma cell–dominated stromal patterns (SG 0) with decreased CPS, and suggest that immune-infiltrated TLS variants (SG 2) are characteristic for adenocarcinoma, underscoring the relevance of spatial immune architecture in lung cancer.

Next, we used PLDA.75 cellular neighborhoods for analysis of tumor slides in the cohort. We clustered the tumor slides into 11 distinct SGs and associated them with clinical parameters and survival data (Fig. 5E-5H, Extended Fig. 5C,D, Extended Table 7). As three tumor slides were collected per patient, we introduced a *heterogeneity measure* (HG) quantifying how many different SGs a patient’s slides were assigned to (Fig. 5G). Such defined spatial heterogeneity emerged as a powerful predictor of patient outcome: Cox proportional hazard analysis revealed that patients with maximum heterogeneity (HG 3) showed significantly worse prognosis compared to patients with homogeneous tumor presentations.

The PLDA.75 analysis again provided higher resolution spatial phenotyping than PLDA.1, and subcategorized some of PLDA.1 groups into distinct biological entities (Fig. 5I). Specifically, SGs 3 and 10 (representative slides in Extended Fig. 5E,F), both characterized by high Ki67 expression, showed distinct immune profiles. SG 3 exhibited greater infiltration of immune cell neighborhoods (CN 20, CN 21) and plasma cell-enriched CN 5, whereas SG 10 contained mostly pure tumor and mural CNs. This aligns with PLDA.1 analysis, where SG 3 corresponded to immune-infiltrated PLDA.1 SG 1, and SG 10 to tumor-immune border PLDA.1 SG 2 (Fig. 5I). Both SG 3 and 10 showed enrichment in ‘other’ (non-adeno and non-squamus) histologies. Additionally, SG 3 was enriched in high CPS and TPS. These results demonstrate that based on their spatial features, proliferative tumor phenotypes separated into divergent, immune-activated versus immune-excluded states with different clinical outcomes. SG 8 was dominated by hypoxic pure tumor CN 10 (Extended Fig. 5G). SGs 5 and 7 both showed high CD15 expression but differed in cellular distribution: SG 7 (representative slide in Extended Fig. 5H) displayed higher neutrophil abundance, while SG 5 (Fig. 5O) showed CD15 enrichment in tumor or near-tumor extracellular areas, suggesting NETosis and neutrophil trapping. SG 5 mapped to non-infiltrated PLDA.1 SGs 0, while SG 7 corresponded to immune-infiltrated PLDA.1 SG 1 (Fig. 5I). SG 5 patients showed the highest hazard ratio (Fig. 5H), suggesting netosis as a potential prognosis marker. In addition to SG 5, SGs 4 and 6 also represented subcategories of cold tumor PLDA.1 SG 0 (Fig. 5I). SG 4 included slides with either low marker expression (CN 13) or increased PDL1 expression (CN 17) in tumor cells (Fig. 5P), and was associated with low CPS. SG 6, characterized by highest mural-enriched CN 7 abundance (Fig. 5N), showed the second-highest hazard ratio (Fig. 5H). SGs 0 showed tumor-predominant CN 21 with memory T-cell infiltration CN 20, was associated with high CPS, and strongly overlapped with PLDA.1 SG 2. Finally, SG 9 encompassed slides with unusual TCF7-expressing tumor cells (Fig. 5Q) and was associated with low TPS and adenocarcinoma. These fine-grained subtypes reveal how different immune cell contexts—plasma cells, neutrophils, or TCF7+ tumor cells—map to distinct clinical features and outcomes within immune-infiltrated tumors.

### Full resolution analysis of the NSCLC dataset Panel 2 introduces highest obtainable detail of cellular neighbourhoods characterization

To gain maximum granularity level insights from 566 tumor microenvironment-oriented Panel 2 slides from 190 IMMUcan patients we used the full-resolution PLDA.100 (Extended Fig. 6A,B) cell embeddings. As expected, we obtained the highest number of 34 distinct cellular neighborhoods that exposed the finest level of detail about observable spatial patterns in the data. Upon inspecting their cell type composition, we grouped the CNs into the following meta-categories: *pure tumor* (tumor cell-dominated), *infiltrated* (mixed tumor-immune), *CD20+* (B cell-enriched), *macrophage-enriched*, and *PMN-MDSC-enriched* (Fig. 6A,B, Extended Table 8).

**Figure 6.**
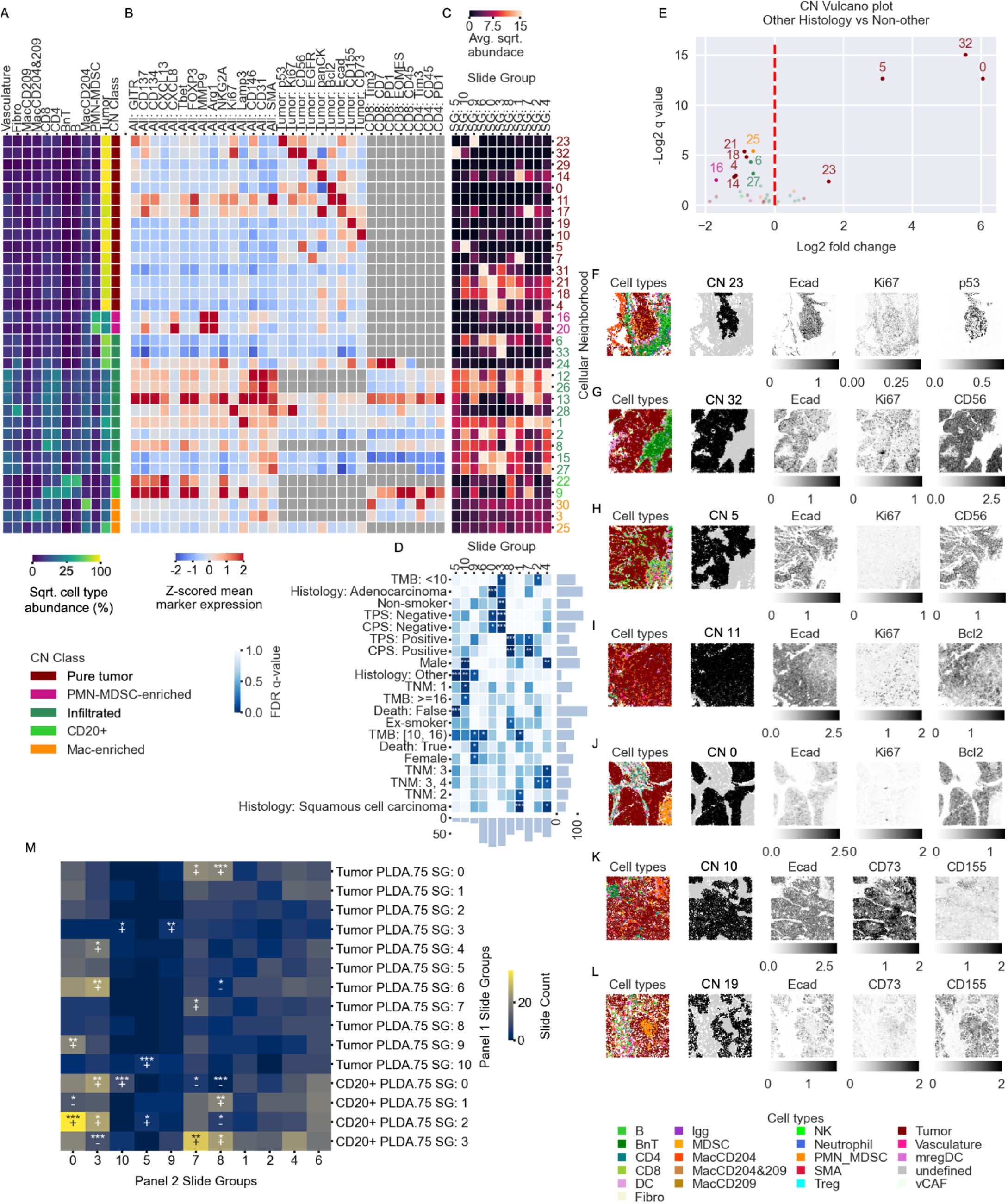
Cellohood Results on IMMUcan NSCLC Panel 2 dataset for PLDA.100 analysis. (**A**) Average cell types (rows) percentage for each CN (columns). (**B**) Z-scored average marker expressions across specific or all cell types (rows) for CNs (columns), presented only for cell types comprising >10% of a given cluster. (**C**-**D**) Slide groups (SGs; columns), characterized by square rooted abundance of CNs (rows) (**C**-**D**) and their association with clinical covariates (**D**). (**E**) Volcano plot of Cellular Neighborhood counts vs ‘other’ histology with log2 fold change on the x-axis and -log2 q value of FDR corrected two-sided t-test on the y-axis. (**F**-**L**) Representative images of slides abundant in a subset of cellular neighborhoods from PLDA.100 analysis. (**M**) Cross-counts of PLDA.100 Panel 2 slide groups with Panel 1 PLDA.75 and CD20^+^ and tumor slide groups with FDR-corrected q-values from two-sided hypergeometric tests (∗q≤0.2, ∗∗q≤0.05, ∗∗∗q≤0.01, + indicating enrichment, - indicating avoidance).

Within pure tumor neighborhoods, we found 15 phenotypically distinct CNs. CNs 23 and 32 showed elevated Ki67/CD56 expression, with CN 23 additionally expressing p53 (representative slides in Fig. 6F,G). CN 5 also exhibited CD56 abundance (Fig. 6H). CNs 29 and 7 consisted of EGFR+ tumor cells, with CN 29 co-expressing pan-cytokeratin. Pan-cytokeratin also characterized CNs 14 and 17, the latter with E-cadherin upregulation. BCL2 upregulation distinguished CNs 0 and 11, with CN 11 showing E-cadherin co-expression (Fig. 6I,J). Immune evasion markers CD155 and CD73 were preferentially expressed in CNs 19 and 10, respectively (Fig. 6K,L). CNs 31, 21, 18, and 4 showed lower marker expression, differentiated by variable E-cadherin/pan-cytokeratin levels. PMN-MDSC-enriched CNs 16 and 20 featured CXCL8-rich environments with Arg1 and MMP9 production. Infiltrated CNs displayed marked heterogeneity, containing T/B cells alongside vascular markers (CD31, CD146, SMA), indicating perivascular immune distribution. The most densely T cell-infiltrated CNs (1, 8, 12, 13, 24, 26) differed in their composition: CN 24 contained CD8+ T cells with elevated CD7 and exhaustion features (PD-1, TIM-3), alongside activation markers (EOMES, CD134, CD137) with varying expression, plus regulatory T cell markers (GITR, FOXP3) and NK inhibitory molecules (NKG2A that pair with CD94 to form an inhibitory receptor). CN 13 displayed a broad marker activation profile, with a similar CD8+ T cell exhaustion phenotype, while CD4+ T cell population showed PD1 and TIM-3 upregulation countered by co-stimulatory CD134 (OX40) expression. For both of these CNs, the presence of EOMES, Tbet, and FOXP3 transcription factors with mixed co-stimulatory and checkpoint molecules indicates short-lived effector T cells in mixed activation/exhaustion states countered by infiltrated Tregs. Additionally, CNs 1, 12, 13 and 26 included myeloid-origin antigen-presenting cells producing Arg1, which impairs T cell activity by downregulating the CD3ζ chain and disrupting TCR signaling. Finally, CN 28 featured the highest levels of proliferating (Ki67+) fibroblasts, alongside activated/exhausted immune cells.

CD20+ cellular neighborhoods (CNs 9 and 22) both exhibited high CXCL13 levels, a strong B-cell chemoattractant, but showed distinct cellular compositions. CN 22 was predominantly populated by B cells, possibly forming follicle-like TLS structures. In contrast, CN 9 displayed mixed B and T cells characteristic of extrafollicular TLS regions. T cells within CN 9 were activated, evidenced by increased TIM-3, CD134, CD137, and CD45 expression, with CD8+ T cells expressing EOMES transcription factor—necessary for full effector differentiation and memory formation but also implicated in exhaustion. Accordingly, T cells in CN 9 characteristic of memory or effector phenotypes expressed high exhaustion marker levels. Additionally, CN 9 contained Treg markers (GITR, FOXP3) and the NK cell negative regulator NKG2A.

Finally, we applied the clinical association pipeline to PLDA.100-derived cellular neighborhoods from the Panel 2 specimens. Clustering analysis revealed 11 distinct slide groups with specific clinical associations (Fig. 6C,D, Extended Fig. 6C,D). SGs 5 and 10, characterized by predominance of CNs 32 (Ki67+, CD56+) and 5 (CD56+), respectively, were significantly overrepresented in rare histological subtypes (“other” histology). Indeed, CNs 32 and 5 were among the most overexpressed in rare histologies (Fig. 6E). Between these SGs, outcomes varied markedly. All SG 5 patients survived (Extended Fig. 6C), indicating that proliferative CD56+ tumors may correlate with better prognosis (Extended Fig. 6E). SG 10 featured male predominance, suggesting sex-specific differences. These groups corresponded to Ki67+ Panel 1 PLDA.75 SGs, with Panel 2 SG 5 mapping to Panel 1 Tumor SG 10, and Panel 2 SG 10 to Panel 1 CD20+ SG 0 (Fig. 6M). SG 0 featured pure tumor CNs 21, 18, and 4 with modest E-cadherin/pan-cytokeratin expression and was enriched in adenocarcinomas. SG 3 showed infiltrated CNs 6, 15, 27 and 33, where tumor cells interfaced with immune cells lacking characteristic activation markers. SG 3 frequently presented low CPS/TPS and non-smoking status. In contrast, SG 8, enriched for CD20+ infiltration in CNs 2, 8, 12 and 26, was associated with elevated CPS/TPS. SG 7 was dominated by CNs 12, 13, and 26, together with the B-cell–rich CN 9, and also showed elevated CPS. SG 1, defined by EGFR+ CNs, was linked to squamous histology. These findings underscore that distinct tumor neighborhoods resolved in full resolution show large heterogeneity of tumor microenvironment and map to clinically divergent lung cancer phenotypes.

To demonstrate that Cellohood generalizes in its ability of establishing detailed cellular neighbourhoods beyond lung cancer, we applied it to an IMC breast cancer dataset of Jackson *et al.* [**6**] at full resolution (PLDA.100), identifying 27 distinct cellular neighborhoods spanning tumor, stromal, and immune microenvironments (Extended Text and Extended Fig. 7,8). Slide-level CN compositions stratified tumors into clinically relevant subgroups beyond receptor status and grade, linking specific proliferative, basal-like, and immune-infiltrated CNs to patient outcomes. Notably, tumor–stroma interface neighborhoods emerged as key determinants of poor prognosis, even with immune infiltration, suggesting breast cancer subgroups that transcend standard hormone receptor/HER2 categories.

## Discussion

Here, we introduced Cellohood, a permutation-invariant set transformer auto-encoder for the identification and analysis of CNs. By preserving the complete molecular signal, our model overcomes limitations of existing spatial analysis methods that rely on aggregated or histogram-based representations and thereby compress biologically relevant heterogeneity. Likewise, while most available methods operate at a fixed resolution or provide limited interpretability, Cellohood enables multi-resolution analyses and produces embeddings that can be directly mapped back to spatial features. Finally, Cellohood integrates clinical association testing, thereby connecting spatial tissue organization directly to patient-level outcomes.

In application to the Visium 10X human DLPFC dataset [**4**], Cellohood uncovered latent topologies aligned with cortical layers, capturing biologically coherent transitions within the tissue. In the CODEX murine spleen model [**5**], it recapitulated known anatomical compartments while distinguishing lupus stage-specific CNs, revealing subtle shifts in B- and T-cell composition, activation states, and spatial localization. For the IMMUcan NSCLC cohort, minimal resolution analysis retrieved CNs on the expected immune-cold to immune-hot spectrum, while standard resolution (PLDA.75) refined these into subclasses, capturing proliferative, hypoxic, PD-L1–positive, and immune-enriched neighborhoods as well as neutrophil trapping phenotypes, and linking intrapatient spatial tumor heterogeneity to prognosis. At maximal resolution (PLDA.100), Cellohood further subclassified rare histological subtypes into biologically coherent groups defined by proliferative (Ki-67–high), p53-aberrant, and CD56-enriched neighborhoods, revealing subtypes that are not captured by conventional histopathology. In the breast cancer cohort [**6**], Cellohood maximal resolution analysis again enabled CN-based subclassification, identifying prognostically distinct subgroups, including immune-enriched, basal-like, and tumor–stroma interface neighborhoods, which stratified patients within standard categories. Taken together, across transcriptomic and proteomic platforms, mouse and human tissues, and healthy and diseased states, Cellohood consistently revealed biologically and clinically relevant CNs, providing a comprehensive framework for characterizing tissue organization in health and disease.

While Cellohood addresses key limitations of previous neighborhood modeling approaches, it still has constraints that motivate further development. First, the current implementation treats all cells within a bag equally, whereas it may be important to account for spatial subcomponents such as the central cell, immediate neighbors, and successive layers of neighboring cells. Second, the analysis focuses on individual cell bags and only implicitly captures their higher-order organization into spatially coherent CNs through clustering. Third, all currently available datasets—including those analyzed here—cover only limited regions of tissue, which may reduce the statistical power of the clinical association pipeline. Despite these limitations, Cellohood demonstrated strong performance in transforming spatial molecular data into biological and clinical insights. As spatial profiling technologies become more accessible and clinically integrated, we anticipate that Cellohood will not only accelerate mechanistic discoveries in tissue biology and immunology, but also inform the development of spatial biomarkers and therapeutic strategies tailored to the microenvironmental context of disease.

## Online methods

### Cellohood Analysis

The Cellohood analysis pipeline proceeds in the following steps:

i. Data preprocessing
ii. Detection of cell (spot) bags
iii. Training of the Cellohood model
iv. Local smoothing of cell and bag embeddings
v. Clustering of cell or bag embeddings into cellular neighborhoods
vi. Multi-resolution principal latent dimension analysis
vii. Clustering of slide representations based on cellular neighborhoods’ abundance
viii. Statistical analysis of association between cellular neighborhoods and clinical features. The pipeline has three user-defined hyperparameters, influencing the granularity of the analysis: First, the user decides whether to operate on cell (spot) or bag embeddings in step
ix. (v). Second, the user picks the resolution by selecting the number of principal latent dimensions in step (vi). Finally, the user picks one of the selected cluster numbers in step (vii).

### Data preprocessing

#### DLPFC Visium 10x [4]

For the analysis of this dataset, we normalized total gene counts to 10^6^, applied log1p transformation and subsequently selected the top 5000 highly variable genes. We then employed SCVI [**25**] with 5-dimensional latent space to embed the gene expression profiles and mitigate batch effects. The dataset consisted of 47681 spots from 12 Visium10x slides (coming from 3 patients, 4 slides each). Each spot was classified to either one of 6 cortical layers (Layers 1 to 6) or white matter (WM).

#### CODEX Mouse Spleen Data [5]

For the analysis of CODEX Mouse Spleen dataset [**5**], in order to eliminate batch effects and normalize marker expression values, we applied the Box Cox quantile transformation for each of the 9 samples separately. This transformation adjusted the distribution of each marker to follow the Gaussian distribution. We used the following list of markers: CD45, Ly6C, TCR, Ly6G, CD19, CD169, CD106, CD3, CD1632, CD8a, CD90, F480, CD11c, Ter119, CD11b, IgD, CD27, CD5, CD79b, CD71, CD31, CD4, IgM, B220, ERTR7, CD35, CD2135, CD44, NKp46, MHCII. Cells were extracted from 9 mice samples: 3 healthy: BALBc-1, BALBc-2, BALBc-3, 3 early lupus MRL-4, MRL-5, MRL-6, 2 intermediate lupus: MRL-7, MRL-8, 1 late lupus MRL-9. 734103 cells were divided into the following cell types: B cells, B220(+) DN T cells, CD106(+)CD16/32(+)CD31(+) stroma, CD106(+)CD16/32(+)CD31(-)Ly6C(-) stroma, CD106(+)CD16/32(-)Ly6C(+)CD31(+), CD106(-)CD16/32(+)Ly6C(+)CD31(-), CD106(-)CD16/32(-)Ly6C(+)CD31(+) stroma, CD11c(+) B cells, CD3(+) other markers (-), CD31(hi) vascular, CD4(+) T cells, CD4(+)CD8(-)cDC, CD4(+)MHCII(+), CD4(-)CD8(+)cDC, CD4(-)CD8(-) cDC, CD8(+) T cells, ERTR7(+) stroma, F4/80(+) mphs, FDCs, NK cells, capsule, dirt, erythroblasts, granulocytes, marginal zone mphs, megakaryocytes, noid, plasma cells.

### IMC datasets

Both Jackson et al. breast cancer. [**6**] and IMMUcan NSCLC IMC data were normalized using the standard arcsinh(x / 5) transformation (regularly applied to CyTOF data).

### IMMUcan datasets (NSCLC Panel 1 and 2)

NSCLC Panel 1 dataset contained2088390 cells from 741 slides (1495338 cells from 569 tumor slides and 593052 cells from 172 CD20+ slides). They were grouped into the following cell types: immune: B, BnT, CD4, CD8, DC, HLADR, MacCD163, NK, Neutrophil, Treg, pDC, plasma, mural / stromal: Mural, tumor: tumor and undefined.

NSCLC Panel 2 dataset contained 1656630 cells from 566 slides. They were grouped into the following cell types: B, BnT, CD4, CD8, DC, Fibro, Igg, MDSC, MacCD204, MacCD204_CD209, MacCD209, NK, Neutrophil, PMN_MDSC, SMA, Treg, Tumor, Vasculature, mregDC, undefined, vCAF.

### Jackson et al. [6] dataset

Jackson et al. breast cancer data contained 855668 cells from 288 slides from the Basel subcohort. Each cell was assigned with one of the cell type classes: immune, tumor, stroma, vessel or unknown. For analysis we used the following markers: EGFR, ECad, DNA1, DNA2, Estrogen, GATA3, H3-1, Ki67, SMA, Vimentin, cl.PARP, HER2, p53, panCK, CK 19, PR, c-Myc, Fibronectin, Keratin-14, Slug, CD20, vWF, H3-2, CK-5, CD44, CD45, CD68, CD3epsi, CAIX, CK 8/18, CK 7, Twist, p-Histone, S6, mTOR.

#### Detection of cell (spot) bags

Cellohood analyses *cell bags*, i.e. groups of cells, possibly of different types, which are spatially close to each other in the tissue. For spot-based technologies, such as 10x Visum spatial transcriptomics, we analyze *spot bags* (groups of colocalized spots) instead.

We identified cell (spot) bags by applying hierarchical clustering to cell (spot) positions for each single image in every dataset. We used a Ward linkage function with a dataset-specific distance threshold. For all of the datasets analyzed, except for DLPFC Visum10x [**4**], we tested different distance thresholds from a set {25, 50, 75, 100} and selected the ones for which the average neighborhood size was the closest to 5. For DLPFC, due to the regular grid-like structure with equal distances between the spots, we fixed the distance threshold to be 5. Hierarchical clustering was performed using the scikit-learn [**26**] package (version 0.24.2).

#### Cellohood auto-encoder architecture

Cellohood is a permutation-invariant set transformer auto-encoder that accepts as an input a given cell bag representation, in the form of a matrix of marker expressions (columns) for each cell in the bag (rows). The model is designed to flexibly handle cell bags of varying sizes, allowing it to process inputs containing different numbers of cells. For spot-based data, the input representation consists of gene expression values for each spot in the input bag. The model creates latent representations of the input data, which comprise both the individual *cell embeddings* (or *spot embeddings*) and the whole *bag embeddings*. The model is trained to reconstruct the input from the bag embeddings using the decoder.

Both encoder and decoder parts have the standard transformer architecture [**27**], with three transformer layers, each with the feed forward layer of size 256 and with 4 attention heads. The size of the first two layers of the encoder and decoder is 128. The final encoder layer is of size d_l_ = 64, the same as the size of Cellohood latent. The final decoder layer has the size of the number of markers used in an analyzed dataset. Additionally, to match the dimensionality of the first two transformer decoder layers (128), we introduced an additional linear projection layer before the decoder that transforms all the embedding sequence elements to a 128-dimensional space.

The input representation of each cell bag is fed to the encoder transformer layers, producing a new cell embedding vector of size d_l_ = 64 for each cell in the bag. In order to obtain the whole bag embedding, we compute the average of the cell embeddings, resulting in the bag embedding vector of size d_l_ = 64. This bag embedding is transformed back to a 128 dimensional space using the fully connected layer to match the dimensionality of the first two decoder layers. Next, it is copied as many times as the number of cells in the input bag representation and a learnable positional encoding is added to each element of the obtained sequence. Next, the sequence is fed to the decoder, which produces a matrix of predicted marker expressions that are trained to reproduce the original marker expressions of the cells in the input bag.

#### Achieving the permutation invariance of the cellular neighborhood representation

As we have not applied any positional encodings to the encoder input, the Cellohood encoder achieves permutation equivariance of the cell embeddings concerning their input order by using a permutation-equivariance property of the vanilla transformer layer [**27**]. The bag embedding is also permutationally equivariant w.r.t. to the input cell order, because the average function used to obtain the bag embeddings from cell embeddings is permutation-invariant. However, this fact introduces a severe obstacle in training Cellohood as an auto-encoder. Namely, without the order of cells preserved between the input and the output representation, the standard auto-encoder reconstruction loss cannot be computed.

In order to solve this problem, we follow the idea introduced by Winter *et al.* [**28**], by incorporating a separate permutation matching model *PM*, which aims at matching the order of the input and the output of a decoder. This model takes the cell embeddings as the input and produces a scalar score for each cell. Later, we apply the soft-sort [**29**] operator that produces the approximation of the permutation matrix. We later apply this permutation to the positional embeddings added to the decoder input and train this model jointly with the encoder and decoder. Once all models are trained, the *PM* model is learned to match the permutation of the input to the Cellohood model with the output of the Cellohood decoder.

#### Cellohood model training

All the Cellohood models are implemented using Keras (in version 2.7.0) [**30**] with a Tensorflow backend. We use a mean-squared error as a reconstruction loss function to compare the z-scored marker expression values between the input and output cell bag representations. The marker expressions are standard scaled based on each marker’s mean and standard deviation statistics, estimated at the cell level on all cells from the training set. For each dataset, we train Cellohood for 1000 epochs and a batch size of 512 (except for the DLPFC Visium10x [**4**] dataset, for which the number of spot bags was relatively smaller compared to the number of cell bags in the other datasets, so we used a batch size of 256 to increase the number of the training iterations). As an optimizer, we use Adam [**31**] with a learning rate of 10^−4^ and a gradient clipping norm of 1.0.

#### Local Smoothing of Cell and Bag Embeddings for Robust Embedding

During cell bag detection, cells were partitioned into disjoint sets of colocalized cells, which could potentially introduce systematic bias as different partitioning strategies might yield different bag compositions. To address this concern, we refined our embedding approach by applying a local smoothing technique. Specifically, for each cell or bag, we computed its smoothed embedding by averaging the embeddings of all its neighbors within a given radius, equal to the distance threshold that was originally employed to cluster cells into disjoint bags. This smoothing mechanism effectively reduces the sensitivity of our embeddings to the specific partitioning scheme used during preprocessing, resulting in more robust and consistent embeddings that better capture the underlying biological relationships between cells.

#### Multi-resolution principal latent dimension analysis

This multi-resolution capability is achieved by applying Principal Component Analysis (PCA) to the complete set of cell (spot) or bag embeddings from a given dataset. The procedure effectively replaces the original embeddings with vectors obtained from a subset of the principal components. To emphasize that these components are derived from latent embeddings, we refer to each as a *principal latent dimension* (PLD) and to the overall procedure as *principal latent dimension analysis* (PLDA).

We defined four distinct resolution levels to enable analyses across different scales of complexity:

● **Minimal Resolution (PLD.1):** Uses only the first principal latent dimension, capturing the most dominant pattern of variation in the data.
● **Coarse Resolution (PLD.50):** Includes the number of principal latent dimensions that together explain 50% of the total variance, providing a broad overview of major cellular patterns while filtering out finer details. If the number of components

explaining 50% of the variance equals one (i.e., identical to PLD.1), we set the number of components analyzed to two.

● **Standard Resolution (PLD.75):** Utilizes the principal latent dimensions explaining 75% of the total variance, offering a balanced representation that preserves most significant biological signals while maintaining manageable computational complexity. If the number of components explaining 75% of the variance is less than or equal to the number used for PLD.50, we set the number of components analyzed to one greater than that used for PLD.50.
● **Full Resolution (PLD.100):** Retains 100% of the variance from the original embeddings, preserving complete information for analyses that require maximum detail.

For each resolution level (except Minimal), we determined the optimal number of dimensions by selecting the point at which the absolute difference between the target explained variance ratio (50%, 75%, or 100%) and the actual explained variance ratio was minimized. This approach ensures that each resolution level contains precisely the amount of information intended for its respective analytical scale.

#### Clustering cell embeddings into cellular neighborhoods

Having the trained Cellohood model, we retrieve the cell (or spot) embeddings for each input bag, as well as the bag embeddings. Both cell (spot) embeddings, as well as the bag embeddings at a predefined PLDA resolution, are clustered using the KMeans algorithm.

In order to determine the number of clusters we use a Fowlkes–Mallows index-based approach similar to [10]. First we run k independent clusterings with different random seeds, for every number of clusters in a set c_0_, *. . .*, c_n+1_, where for every i = 0, *. . .*, n+1, c_i_ = c_0_+i·step, with step ∈ N_+_. Later, for each i = 1, *. . .*, n, we compute the FM I(i) which is an average Fowlkes–Mallows index between all pairs of different clusterings with numbers of clusters c_i_ and c_i+1_ as well as c_i_ and c_i–1_. This average serves as a measure of the stability of the given number of clusters c_i_ when compared with clusterings with numbers of clusters c_i–1_ and c_i+1_. We later analyze those clustering candidates with c_j_ clusters, for which FM I(j) is a local maximum. From the k clusterings with c_j_ clusters we select as the final clustering the one with the least inertia. The final clustering of cells (spots) or bags defines the cellular neighborhoods.

For the datasets analyzed in this study, we fixed k = 5, c_0_= 5, n = 50, and step = 1.

#### Diffusion Dimensionality Reduction

For the Diffusion Dimensionality Reduction we used a pydiffmap [31] package with Gaussian kernel with ɛ=0.01, and diffusion map parameters set to α=0.1, and k=64.

### Clinical Association Pipeline

#### Cellular Neighborhood Summarization

To summarize cellular neighborhoods, we computed cell type/cell class statistics as well as marker statistics for each CN. For cell type analysis, both at the cell and bag level, we identified all cells assigned to a specific CN and calculated the percentage of each cell type within this set. For marker analysis, we computed statistics either at the level of all cells or at the specific cell type level. For general statistics (denoted as "All"), we calculated the mean marker expression of all cells assigned to a particular CN. For cell-level analysis, we calculated the mean marker expression of all cells assigned to a particular cell type. Subsequently, these statistics were z-scored (standardized) across all CNs to facilitate comparison. Density analysis was performed by computing the mean size of the bag assigned to a particular CN. As all the bags were created to have roughly the same radius, the size of the bag correlates with a density. These mean sizes were once again z-scored across all CNs.

#### Slide Level Aggregation and Clustering into Slide Groups

The identified cellular neighborhoods are mapped onto slides by assigning each cell or spot on the slide to the cellular neighborhood that this cell or spot belongs to. Such slide mapping results in spatially coherent subregions of the slides assigned to their cellular neighborhoods. Having the slide mapping, we determine the numbers of cells (spots) assigned to each cellular neighborhood per slide. For each dataset, we collected the counts of the Cellohood cellular neighborhoods at the slide level in order to obtain slide representations. Due to a small sample size (only 9 mice), we used raw counts to analyze the CODEX Mouse Spleen dataset [**5**]. For all IMC datasets, we used preprocessed counts and ran the cluster selection algorithm to identify specific slide groups that were subsequently used in the analysis. For the breast cancer IMC dataset [**6**], we applied the log1p area-adjusted transformation introduced by the dataset authors in the original paper. For the NSCLC IMC dataset, due to the much greater heterogeneity of the tumor, we applied a square root transformation to raw cellular neighborhood (CN) counts in order to mitigate the effect of outliers.

The obtained slide representations are then clustered using the KMeans algorithm. The number of clusters is determined using the Fowlkes–Mallows index-based approach, as described in Section **Clustering cell embeddings into cellular neighborhoods**.

#### Correlation with clinical data

##### Association of slide groups with clinical covariates

In order to compute associations between slide groups and clinical covariates, we used the following approach: For each slide group and each clinical covariate, we divided patients into those who had at least one slide assigned to the group and those who did not, as well as those who had been assigned a given clinical covariate and those who had not. We then computed a one-sided hypergeometric test to evaluate enrichment of a given clinical covariate for patients in a given slide group. The resulting p-values were further adjusted using the False Discovery Rate (FDR) correction.

##### Cox analysis

For the Cox analysis, we represented each patient as a vector of one-hot indicators denoting whether the patient had a slide from a given slide group. This vector was appended with one-hot indicators of clinical features. Both slide group and clinical feature indicators were z-scored across cohorts to ensure standardization.

##### Slide group cross-count comparison

For cross-count analysis between Immucan NSCLC Panel 1 tumor PLDA.1 and PLDA.75 slide groups, we noted that each slide was assigned to both grouping schemes. Therefore, we performed a two-sided hypergeometric test between each pair of PLDA.1 and PLDA.75 slide groups to assess their association.

For Panel 1 PLDA.75 Tumor and CD20^+^ groups, we used a different approach at the patient level. For each pair of these slide groups, we divided patients into subsets based on whether they had at least one slide assigned to either group. We then performed a two-sided hypergeometric test to evaluate associations at the patient level.

The same comparison methodology was applied for assessing relationships between Panel 2 PLDA.100 and Panel 1 PLDA.75 (both tumor and CD20^+^) slide groups.

##### Clinical data for the IMMUCAN NSCLC cohort

Clinical data for this analysis were obtained as described in the accompanying cohort paper [**15**]. CPS/TPS scores were considered *high* when the percentage of all or tumor-only cells exceeded 1% on the corresponding mIF slide from the same patient. TMB scores below 10 were classified as *low*, those between 10 and 16 as *high*, and those above 16 as *super high*.

##### Heterogeneity measure for IMMUCAN NSCLC slide groups

In this cohort, three tumor slides were collected for each patient. We introduced an additional heterogeneity measure defined as 1 (*low*) if all slides were assigned to the same slide group, 2 (*medium*) if the slides were assigned to two different slide groups, and 3 (*high*) if all three slides were assigned to different slide groups.

### Benchmarking against other methods

#### DLPFC Visium dataset [4]

For this dataset, for both Cellohood and aggregated cell bag (Agg.CBR) representations we adhered to the evaluation protocol and metrics established in the CellCharter paper [**10**]. For the existing baseline methods, we utilized the metric values published in that paper. It is worth noting that we did not include the CytoCommunity method [**11**] in our comparison for the DLPFC Visium dataset [**4**], as this method requires cell/spot type annotations which are not available for this particular dataset.

#### CODEX Mouse Spleen dataset [5]

For this dataset, we followed the evaluation protocol and metrics described in the CytoCommunity paper [**11**]. To ensure fair comparison with existing baseline methods, we used the results published in this same paper.

### Data and Code Availability

Data availability for the NSCLC dataset is outlined in a companion manuscript by Schultz et al. []. Data for Code is available at szczurek-lab github [link]

## Acknowledgements

The IMMUcan project has received funding from the Innovative Medicines Initiative 2 Joint Undertaking under grant agreement no. 821558. This Joint Undertaking receives support from the European Union’s Horizon 2020 Research and Innovation Programme and EFPIA (https://IMI.europa.eu). ESz acknowledges the support from the Polish National Science Centre SONATA BIS grant No. 2020/38/E/NZ2/00305.

## Author Contributions

Conceptualization was carried out by MM, DS, KG, NE, JK, OB, ES and ESz. Software development, visualization and formal analysis were performed by MM, methodology by MM and ESz. Investigation was conducted by DS, NE, OB, ES, JG, HH, DN, MMf and ESz. Resources, specifically IMC data collection, were provided by DS, NE and BB. Data curation was performed by RL, DS, NE and MMf. Writing was performed by MM, with edits by ESz and review by all coauthors. Supervision was provided by ESz. Project administration was carried out by HH, MMf and ESz.

