## Supplementary Material for "Cellohood: multi-granular discovery of cellular neighborhoods with a permutation-invariant set transformer auto-encoder"

#### Cellohood Supplementary

#### Extended Text

##### 1. Simulated CD8 / Macrophages / Tumor bag dataset

To empirically demonstrate the technical challenges related to the full cell bag representation and the limitations of the alternative representations, we generated a simulated dataset of cell bags, each containing exactly three cells expressing five markers: Ecad, CD163, CD8, PD1, and PDL1 (Extended Fig. 1). Marker expression was binary (0 for negative, 1 for positive), with Ecad, CD163, and CD8 used as lineage-defining markers for tumor cells, macrophages, and $CD8^+$ T cells, respectively. We constructed each cell bag to contain precisely one cell of each type, while systematically varying the expression of PD1 and PDL1 to create biologically distinct microenvironments. Specifically, tumor cells were either $PDL1^+$ or $PDL1^-$, $CD8^+$ T cells were either $PD1^+$ or $PD1^-$, and macrophages exhibited three phenotypes: $PD1^+PDL1^-$, $PD1^-PDL1^+$, or $PD1^-PDL1^-$. There resulted in 12 distinct cellular neighborhood types. All of these neighborhoods, by design, were represented by exactly the same histogram cell representations. Additionally the following neighborhoods had exactly the same aggregated cell bag representations:

- *one cell type $PD1^+$ / two cell types $PDL1^-$*: $CD8^+$ $PD1^+$, Tumor $PDL1^-$, Macrophage $PD1^-$ $PDL1^-$; $CD8^+$ $PD1^-$, Tumor $PDL1^-$, Macrophage $PD1^+$ $PDL1^-$,
- *one cell type $PD1^+$ / only one cell type $PDL1^+$* : $CD8^+$ $PD1^+$, Tumor $PDL1^+$, Macrophage $PD1^-PDL1^-$; $CD8^+$ $PD1^-$, Tumor $PDL1^+$, Macrophage $PD1^+PDL1^-$, $CD8^+$ $PD1^+$, Tumor $PDL1^-$, Macrophage $PD1^-PDL1^+$
- *two cell types $PD1^-$ / one cell type $PDL1^+$*: $CD8^+$ $PD1^-$, Tumor $PDL1^+$, Macrophage $PD1^-$ $PDL1^-$; $CD8^+$ $PD1^-$, Tumor $PDL1^-$, Macrophage $PD1^-$ $PDL1^+$.

##### 2. Cellohood PLDA.100 analysis and clinical association pipeline for healthy and lupus affected mouse samples on CODEX Spleen data

For the analysis of the lupus progression, we deployed spatial transcriptomics analysis at full PLDA.100 resolution, identifying 14 distinct cellular neighborhoods (CNs) that revealed characteristic architectural patterns evolving throughout lupus progression (Extended Fig. 3F-I).

Healthy BALBc control samples (BALBc1-3) exhibited continuous and clearly demarcated splenic compartments. The periarteriolar lymphoid sheath (PALS), representing the T-cell zone surrounding central arterioles, was characterized by CN9. B-cell follicles were centered around CN13, while red pulp regions were dominated by CN5. Critically, a prominent marginal zone represented by CN6 formed a distinct barrier separating the white pulp (PALS and B-cell follicles) from the red pulp. This marginal zone organization is essential for filtering blood-borne antigens and mounting rapid immune responses. Additional transitional zones were identified: CN4 and CN0 capturing the outer boundaries of B-cell follicles, and CN7 demarcating the interface between PALS and B-zone compartments, maintaining precise lymphocyte segregation necessary for coordinated immune responses.

Early lupus samples (MRL4-6) demonstrated significant molecular and spatial alterations. The healthy PALS CN9 was replaced by CN2, containing T cells with elevated expression of MHC II, CD44, and CD45 markers, indicating enhanced antigen presentation capacity and an activated phenotype. B-zone centers became dominated by CN3, characterized by exceptionally high expression of CD5, CD79b, CD45, B220, and IgD on B cells. This CD5+ B cell expansion is particularly significant, as these populations are associated with autoantibody production in lupus, though paradoxically may also attempt regulatory functions through IL-10 production.

Marginal zone architecture showed notable disruption, with increased abundance of transitional CNs 7, 4, and 0, but decreased frequency of the primary marginal zone CN6. This alteration likely impairs the spleen's filtering function and clearance of apoptotic debris, potentially contributing to autoantigen exposure. The red pulp CN5 was replaced by CN1, enriched with CD71+Ter119+ erythroblasts, indicating stress erythropoiesis—a compensatory response to chronic inflammation-associated anemia.

Intermediate and late lupus stages exhibited severe architectural disruption. B-zones became heterogeneous, containing both the aforementioned CN3 and CN8—a pathological neighborhood mixing CD5+ B cells with T cells, suggesting loss of normal lymphocyte compartmentalization. Red pulp regions were overtaken by CN10, displaying complex cellular composition: CD27+CD71+Ter119+ erythroblasts indicating ongoing extramedullary hematopoiesis, alongside CD5+CD79b+B220+IgD+ B cells with aberrant red pulp localization.

This neighborhood was enriched in ERTR7 (marking fibroblastic reticular cells), IgM, Ly6G (neutrophils), and Ly6C (inflammatory monocytes), indicating extensive stromal remodeling and inflammatory infiltration. The co-localization of these diverse populations within single neighborhoods represents complete breakdown of normal splenic architecture, transforming organized immune compartments into chaotic inflammatory sites where normal immune regulation cannot occur, perpetuating autoimmune pathology.

#### 1. Clinical association pipeline reveals associations of cellular neighborhoods with clinical features in breast cancer

To showcase applicability of our framework to different cancer types, we also applied Cellohood and the clinical association pipeline to gain novel insights from the IMC breast cancer dataset published by Jackson *et al*. [**1**]. At full resolution (PLDA.100), we identified 27 distinct CNs that captured the diverse tumor microenvironments (Extended Fig. 7A, B, Extended Table 9). These neighborhoods encompassed pure tumor regions, tumor–stroma interfaces, stroma-dominated areas, and immune-rich compartments. Pure tumor neighborhoods showed marked heterogeneity: CN 8 (c-Myc and hypoxia marker CAIX), CN 15 (basal cell markers), CN 26 (luminal markers including PR and CK8/18), CN 9 (Ki67+ proliferating tumor cells), and cytokeratin-rich CN 23 (with multiple epithelial markers). Two HER2-enriched neighborhoods were identified: CN 2 (co-expressing p53, EGFR, cleaved PARP) and CN 12 (ER and GATA3).

At tumor–stroma interfaces, CN 0, 6, and 24 contained higher proportions of tumor cells, with CN 0 showing mTOR activation and CN 24 the highest overall cell density. Immune-cold interfaces included CN 21 and CN 4, enriched in non-classified epithelial cells, i.e., epithelial cells with low expression of the measured markers, and CN 7 (tumor cells expressing poor prognosis markers: c-Myc, Twist, phosphorylated histone). In contrast, CN 17, 16, and 11 formed immune-hot interfaces; CN 16 showed apoptosis in tumor cells and fibroblast proliferation, while CN 11 had high DNA content across cell types. Among stroma-dominated regions, CN 3 was enriched for CAF activation, and CN 19 represented stromal–vessel interfaces. Immune-dominated neighborhoods included CN 1 (macrophages and activated fibroblasts) and CN 13 (activated B and T cells).

Additional Cellohood embedding analysis indicated a clear organization of the latent space of the model by expression of key markers, such as panCK (Extended Fig. 8A), Carbonic Androhydrase (Extended Fig. 8B), Vimentin (Extended Fig. 8C), and CD45 (Extended Fig. 8D), as well as clustering into cell classes (Extended Fig. 8E).

Next, following the clinical association pipeline, we aggregated CN counts per patient slide and clustered them into SGs based on CN composition (Extended Fig. 7C, Extended Fig. 8F,G). Associations with clinical features (Extended Fig. 7D) divided SGs into three groups: HR-negative (SG 5, 2, 10), HR-positive (SG 11, 0, 7, 6), and *clinically non-classifiable*, defined as subgroups lacking strong association with hormone receptor status (SG 3, 8, 4, 1, 14).

In the HR-negative group, SG 5 differed from SG 10 and 2 by enrichment in inflammatory CN 13. SG 5 also contained epithelial-rich CNs 23, 20, and 18, and was the only HR-negative subgroup not enriched in Grade 3 tumors, suggesting distinct biology. SG 10 featured proliferative CN 9 (Ki67+) and reduced CN 5 and 21 abundance (non-classified epithelial cells). Cox analysis showed SG 2 had the lowest hazard ratio (Extended Fig. 7E), and split Grade 3 patients into groups with different survival outcomes (Extended Fig. 7F), highlighting the heterogeneity of high-grade tumors and the limitations of conventional grading systems.

In the HR-positive group, SG 11 and SG 6 were enriched in tumor CK+

CN 23, consistent with luminal epithelial characteristics. SG 6 also had the highest abundance of basal-like CN 15 (CK5+, CK14+). This subgroup displayed the highest hazard ratio among all SGs (Extended Fig. 7E), consistent with the poor prognosis associated with basal marker expression even in HR-positive contexts. Moreover, SG 6 stratified HR+HER2– patients into two subgroups with significantly different survival outcomes (Extended Fig. 7G), underscoring clinically relevant heterogeneity within this common subtype. Spatial arrangement of the CNs per each SG was in agreement with their identified highly expressed markers (Extended Fig. 7H,I).

Together, these analyses demonstrate that distinct CN compositions found by Cellohood stratify breast tumors into clinically relevant subgroups beyond conventional grading and receptor status. Specific neighborhoods, such as proliferative, basal-like, or immune-infiltrated CNs, defined patient survival groups and uncovered heterogeneity within HR-negative, HR-positive, and high-grade tumors. Notably, tumor–stroma interface CNs emerged as key determinants of poor outcome, even in the presence of immune infiltration, highlighting the prognostic value of spatially resolved microenvironmental features.

Slide-level correlation analysis of cellular neighborhood abundances revealed several co-occurrence patterns within the tumor microenvironment (Extended Fig. 8F). Tumor–stroma interface neighborhoods (CN 11, 17, 4, 21, 14, and 22) frequently coexisted within the same tumors. Likewise, tumor-dominated regions expressing ER and GATA3 (CN 10 and 12) showed strong co-occurrence, consistent with their shared luminal phenotype. Correlations were also observed between tumor-dominated regions containing non-classified epithelial cells (CN 5, 18) and neighborhoods with tumor cells lacking high expression of conventional tumor markers (CN 25, 6, 20). These phenotypically ambiguous tumor populations may therefore represent related states or lineages that commonly coexist within the same tumors. CN 13, enriched in activated immune cells, tended to co-occur with CN 7, characterized by high expression of aggressive tumor markers, suggesting an active immune response at sites of tumor progression. Together, these co-occurrence patterns are consistent with biological processes known to take place at the invasive front of cancers.

### Extended Figures

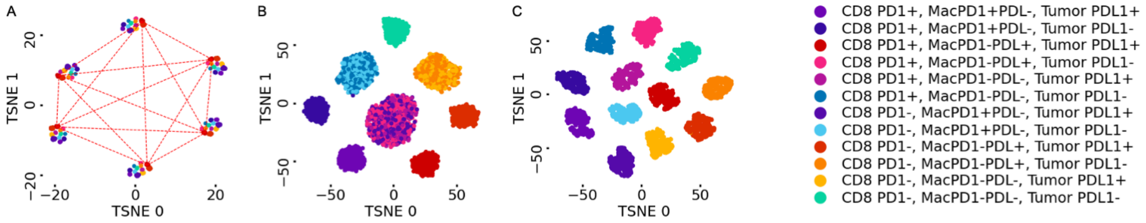


**Extended Figure 1. Simulated Dataset overview**

(A) T-SNE visualization demonstrating the permutation dependence problem. Different permutations of the same cell bag (B1) are represented as distinct objects.

(B) T-SNE visualization demonstrating the information loss of aggregated cell bag representation. Although it solves the permutation dependence problem, it loses cell-specific information.

(C) Cellohood bag embeddings maintain both permutation invariance and cell-specific information.

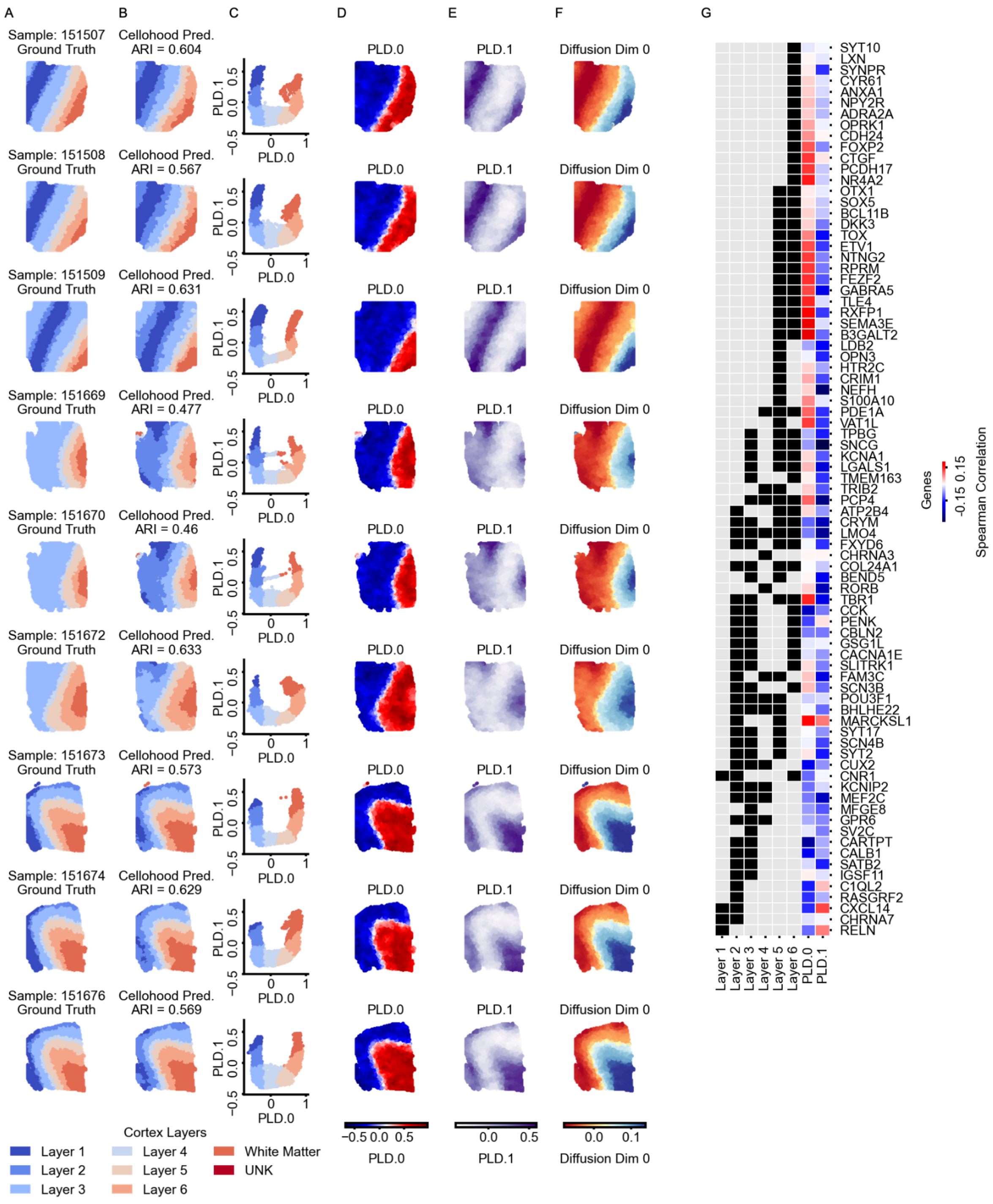


**Extended Figure 2. DLPFC**

**(A–F)** Visualization of Cellohood results for a cortical layer assignment task. Panel (A) shows the ground truth annotation, and (B) shows Cellohood predictions. Panels (C–F) display PLDA.50-based representations, providing insight into the model's internal feature extraction. **(G)** Spearman correlations (color-coded) between the gene expression of cortical layer marker genes from **[2]** and **[3]**, and the PLD.0 and PLD.1 directions. Marker gene status for each layer is indicated by black entries.

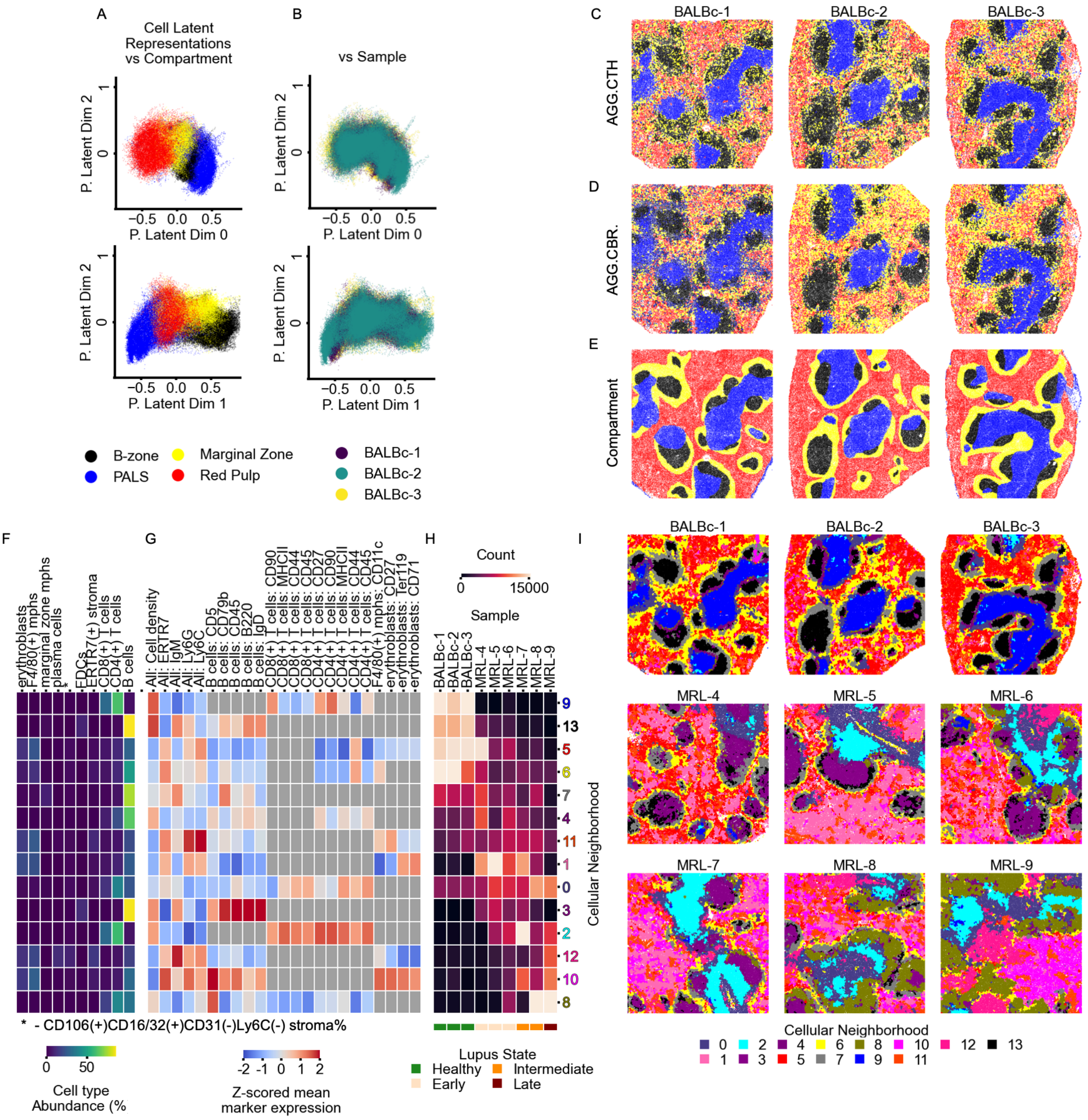


**Extended Figure 3. Cellohood performance and clinical association pipeline for healthy and lupus affected mouse samples on CODEX Spleen data [4]**

(A–B) PLDA.50 representations of cells: separation of splenic regions (B-zone, PALS, Marginal Zone, Red Pulp) (A) and similarity among three healthy controls (BALB/c-1–3) (B). (C–E) Predictions from CTH.CBR (C) and AGG.CBR (D) compared with ground-truth compartment annotations (E). (F-H) Heatmaps of CN characteristics: cell type proportions (F), z-scored marker expression (G), and CN abundance (H). Column colors denote disease state (healthy, early, intermediate, late). (I) Spatial mapping of CNs across nine samples (three healthy BALB/c, six lupus-affected MRL/lpr), identifying splenic compartments across disease states.

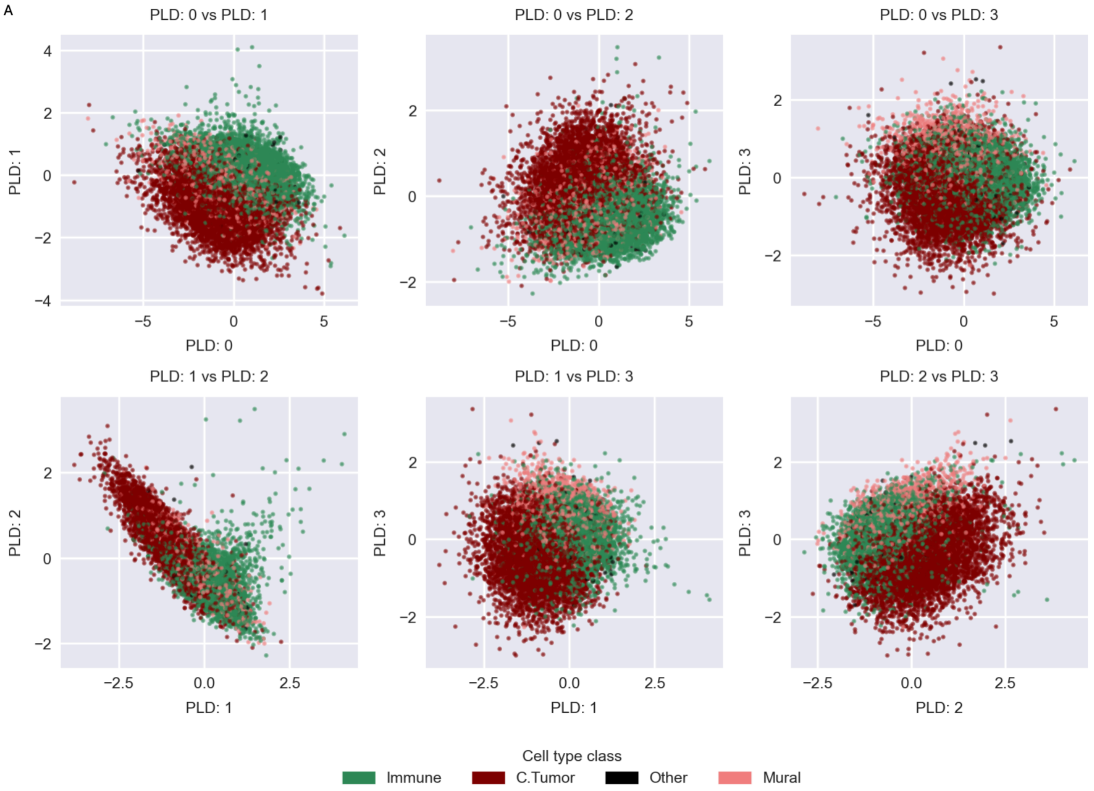


**Extended Figure 4. PLD for NSCLC Panel 1 tumor slides**
(A) Pairs of first 4 Principal Latent Dimensions (PLD) colored by cell type class.

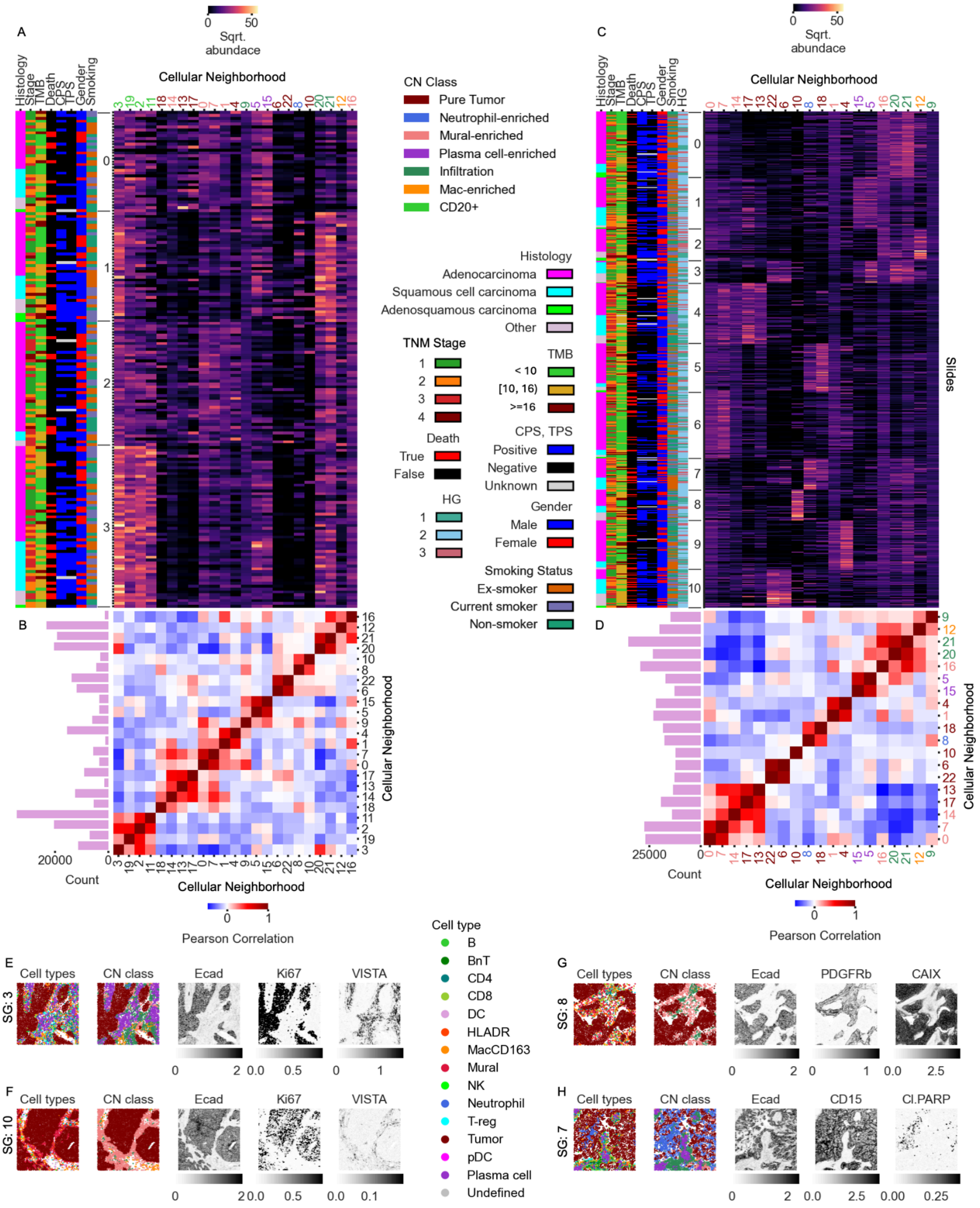

**Extended Figure 5. NSCLC Panel 1**

(A-B) Analysis of the IMMUcan NSCLC Panel 1 tumor slides. (A) Abundances of CNs (columns) for individual slides (rows). Colors in leftmost columns: clinical covariates associated with each patient. (B) Correlation between CN abundances at the slide level. Side bars: total CNs count across the cohort. (C-D) Analysis of the IMMUcan NSCLC Panel 1 CD20+ slides. (C) Abundances of CNs (columns) for individual slides (rows). Colors in leftmost columns: clinical covariates associated with each patient. (D) Correlation between CN abundances at the slide level. Side bars: total CNs count across the cohort. (E-H) Representative slides from SG 3 (E), SG 10 (F) SG 8 (G) and SG 7 (H), with cell types and selected marker expressions.

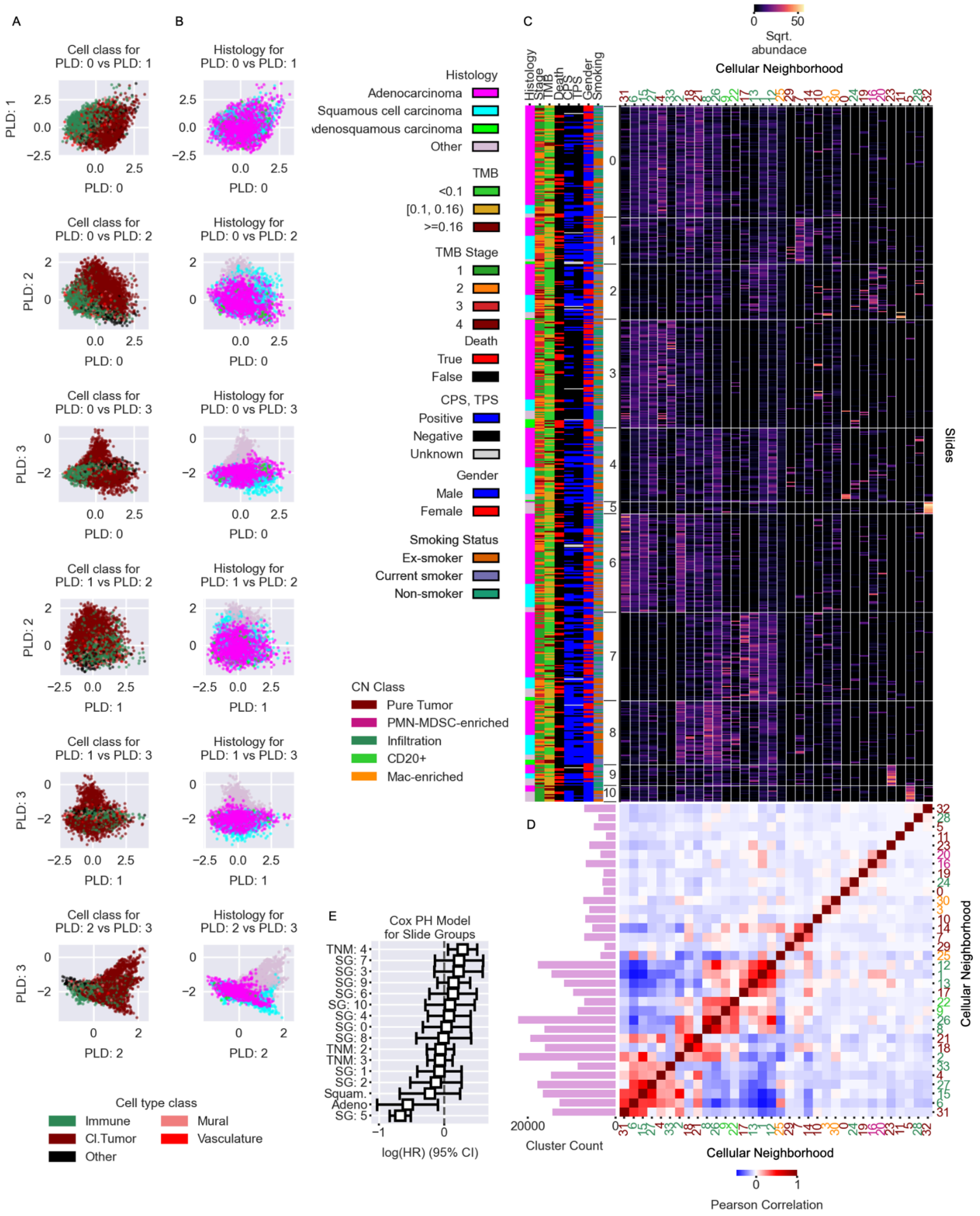


**Extended Figure 6. Immucan NSCLC Panel 2**

(A-B) Pairs of first 4 Principal Latent Dimensions (PLD) colored by Cell type class (A) and histology of patient (B). (C) Abundances of CNs (columns) for individual patients (rows). Colors in leftmost rows: clinical covariates associated with each patient. (D) Correlation between CN abundances at the slide level. Side bars: total CNs count across the cohort. (E) Multivariate Cox proportional hazard model for SGs and relevant clinical covariates, and survival of TNM Stage 1 patients established as the baseline. Error bars represent 95% confidence intervals for each hazard ratio.

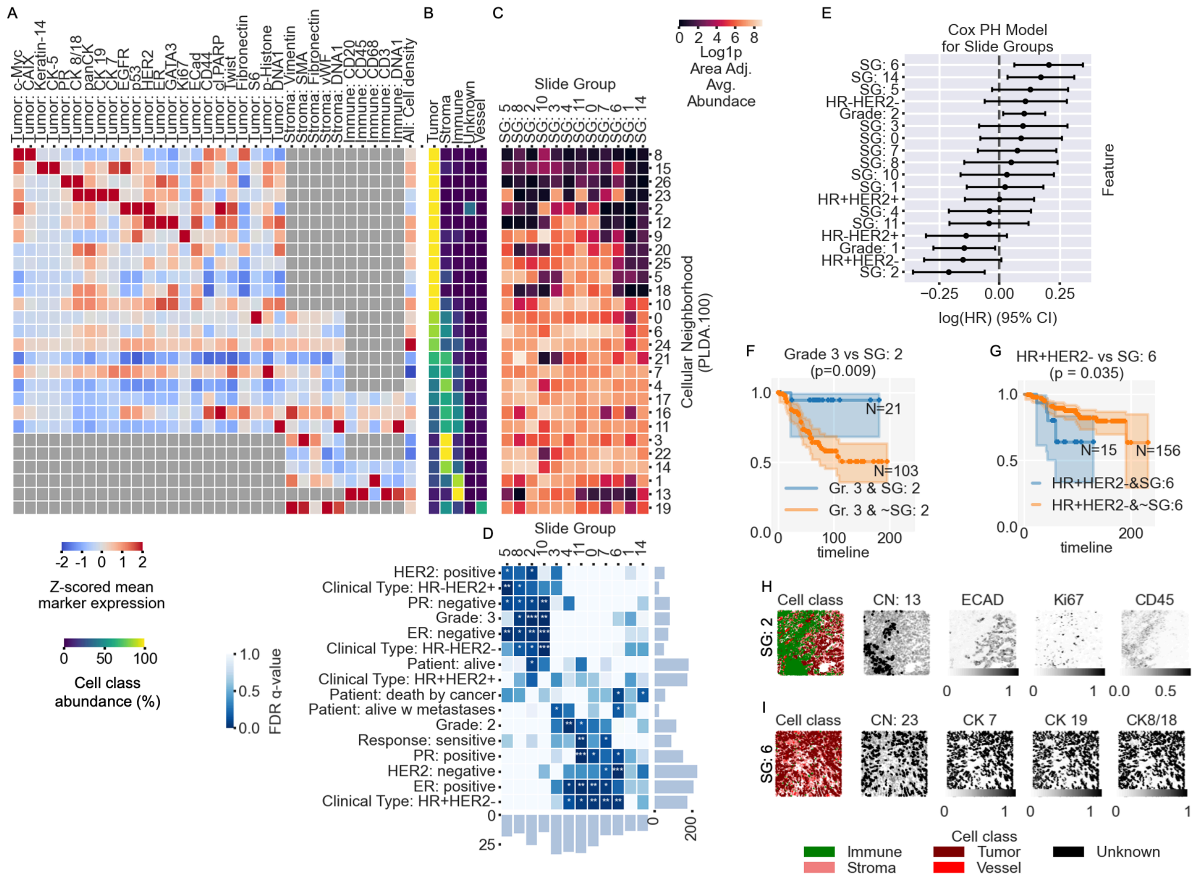


**Extended Figure 7. PLDA.100 cell embedding-based cellular neighborhoods (CNs) and their clinical relevance for Breast Cancer dataset (Jackson et al. [*1*])**

(A) Z-scored average marker expressions across specific or all cell classes (rows) for CNs (columns), presented only for cell classes comprising >10% of a given cluster. (B) Average cell class (rows) percentage for each CN (columns). (C) Slide groups (SGs; columns), characterized by average log1p area adjusted abundance of CNs (rows). (D) Association of SGs (columns) with clinical covariates (columns), with FDR-corrected q-values from one-sided hypergeometric tests performed for each SG against each clinical variable (∗q≤0.2, ∗∗q≤0.05, ∗∗∗q≤0.01). (E) Cox proportional hazard model for SGs and relevant clinical covariates, and survival of Grade 3 patients established as the baseline. Error bars represent 95% confidence intervals for each hazard ratio. (F) Kaplan-Meier survival curve for Grade 3 patients stratified by SG 2. (G) Kaplan-Meier curves for $HR^{+}HER2^{-}$ patients stratified by SG 6. (H-I) Representative slides from SG 2 (H) and SG 6 (I), with cell types and selected marker expressions.

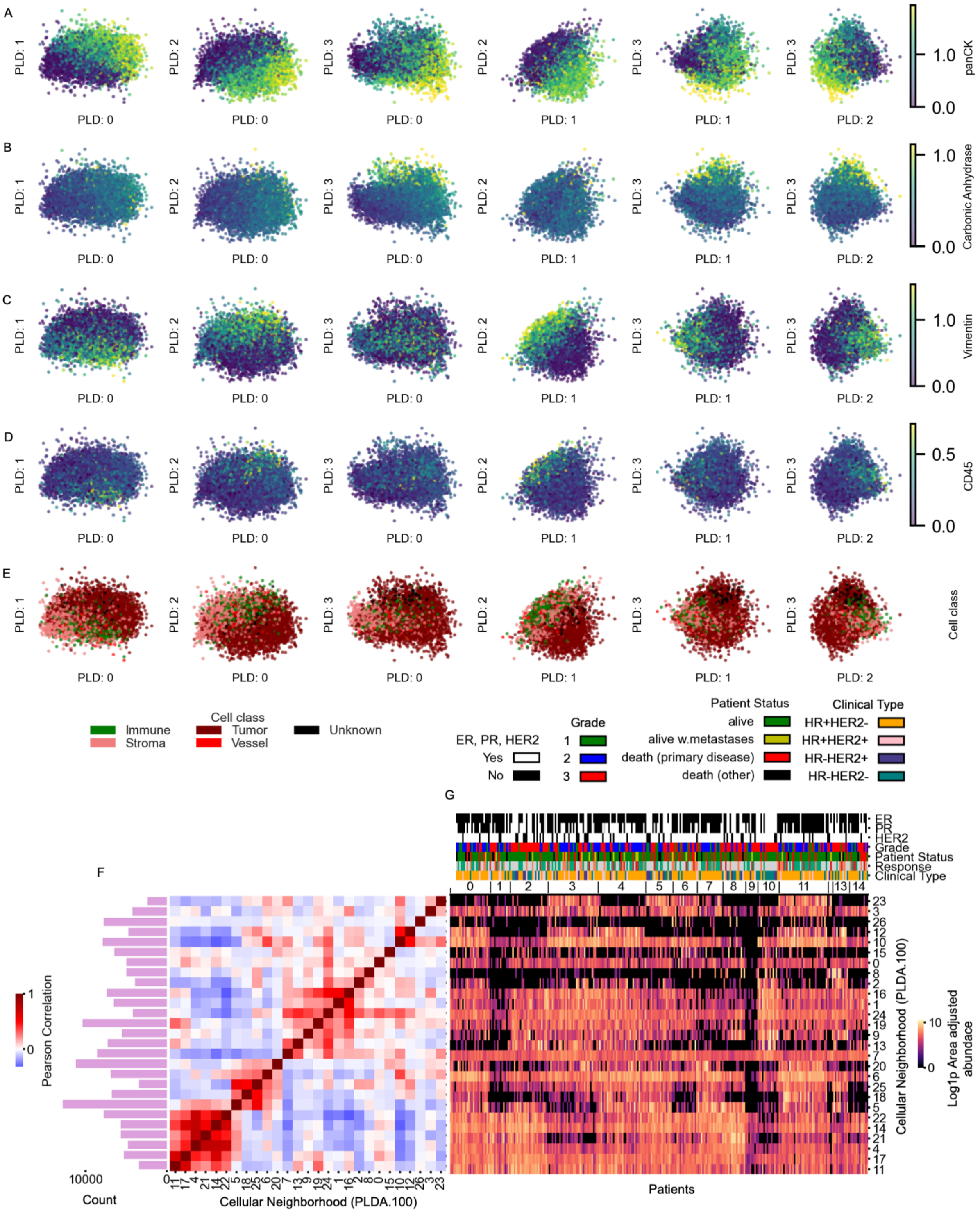


**Extended Figure 8. Cellohood analysis of Jackson et al. 2019**

(A-E) Pairs of first 4 Principal Latent Dimensions (PLD) colored by panCK (A), carbonic anhydrase (B), vimentin (C), CD45 (D) and cell type class (E). (F) Correlation between CN abundances at the slide level. Side bars: total CNs count across the cohort. (G) Abundances of CNs (rows) for individual patients (columns). Colors in top-most rows: clinical covariates associated with each patient

#### Extended Tables

| Marker | Function | Cell Type |
|---|---|---|
| **MPO** | Enzyme producing ROS in neutrophils; antimicrobial defense | Neutrophils |
| **Histone H3** | Nuclear protein, chromatin structure | All nucleated cells (nuclear marker) |
| **SMA (α-Smooth Muscle Actin)** | Contractile protein, cytoskeletal structure | Mural cells (Smooth muscle cells, myofibroblasts, pericytes) |
| **CD16** | Fc receptor mediating ADCC, phagocytosis | NK cells, neutrophils, monocytes |
| **CD38** | Ecto-enzyme, activation marker | Plasma cells, activated T/B cells, NK cells |
| **HLADR** | MHC class II, antigen presentation | Dendritic cells, monocytes, macrophages, activated T cells |
| **CD27** | Co-stimulatory receptor, memory formation | Memory B cells, memory T cells, plasma cells |
| **CD15** | Adhesion molecule, granulocyte marker | Neutrophils, eosinophils |
| **CD45RA** | Isoform of CD45, naïve marker | Naïve T cells |
| **CD163** | Scavenger receptor, anti-inflammatory | Macrophages |
| **B2M** | MHC class I component | All nucleated cells (esp. lymphocytes) |
| **CD20** | B-cell receptor component | Mature B cells |
| **CD68** | Lysosomal glycoprotein, phagocytosis | Macrophages, monocytes |
| **Ido1** | Tryptophan-catabolizing enzyme, immune suppression | DCs, macrophages |
| **CD3** | TCR complex component | Pan-T cells |
| **LAG3** | Inhibitory checkpoint receptor | Exhausted/activated T cells |
| **CD11c** | Integrin αXβ2, adhesion, phagocytosis | Dendritic cells, some macrophages |
| **PD1** | Immune checkpoint receptor | Exhausted/activated T cells, some B cells |
| **PDGFRb** | Growth factor receptor, pericyte biology | Pericytes, fibroblasts |
| **CD7** | T-cell receptor accessory, signaling | T cells, NK cells |
| **GrzB** | Cytotoxic protease | Cytotoxic T cells, NK cells |
| **PDL1** | Immune checkpoint ligand | Tumor cells, macrophages, DCs |
| **TCF7** | Transcription factor, memory stemness | T cells |
| **CD45RO** | Isoform of CD45, memory marker | Memory T cells |

| | | |
|---|---|---|
| **FOXP3** | Transcription factor, regulatory program | Regulatory T cells (Tregs) |
| **ICOS** | Co-stimulatory receptor | Activated T cells |
| **CD8a** | Cytotoxic receptor chain | $CD8^+$ T cells |
| **Carbonic Anhydrase** | pH regulation, hypoxia marker | Hypoxic tumor cells |
| **CD33** | Sialic acid-binding lectin, inhibitory receptor | Myeloid progenitors, monocytes, MDSCs |
| **Ki67** | Nuclear proliferation marker | Cycling cells |
| **VISTA** | Immune checkpoint receptor | Myeloid cells, some T cells |
| **CD40** | Co-stimulatory receptor | B cells, dendritic cells, macrophages |
| **CD4** | TCR co-receptor | $CD4^+$ T helper cells |
| **CD14** | LPS receptor, innate activation | Monocytes, macrophages |
| **Ecad** | Adhesion molecule | Tumor cells |
| **CD303** | C-type lectin receptor | Plasmacytoid dendritic cells (pDCs) |
| **CD206** | Mannose receptor, endocytosis, tissue repair | Macrophages, DCs |
| **cl.PARP** | Apoptosis marker (DNA repair enzyme cleavage) | Apoptotic cells |
| **DNA1** | DNA intercalator (nuclear segmentation) | All nucleated cells |
| **DNA2** | DNA intercalator (nuclear segmentation) | All nucleated cells |

**Extended Table 1: Panel 1 IMC Markers: Function and Cell Type**

| Marker | Function | Cell Type |
|---|---|---|
| **panCK** | Cytoskeletal intermediate filament, epithelial marker | Tumor / Epithelial cells |
| **Histone H3** | Nuclear protein, chromatin structure | All nucleated cells (nuclear marker) |
| **SMA (α-Smooth Muscle Actin)** | Contractile protein, cytoskeletal structure | Mural cells (smooth muscle cells, myofibroblasts, pericytes) |
| **CD7** | T-cell receptor accessory, signaling | T cells, NK cells |
| **CD11b** | Integrin αMβ2, adhesion, migration, phagocytosis | Monocytes, macrophages, granulocytes, NK cells |
| **Arg1** | Enzyme in arginine metabolism, immune suppression | Macrophages, myeloid-derived suppressor cells (MDSCs) |
| **CD146 (MCAM)** | Adhesion molecule | Endothelial cells, pericytes, some T cells |
| **EGFR** | Growth factor receptor, proliferation | Epithelial cells, tumor cells |
| **CD45** | Leukocyte common antigen, pan-immune marker | All hematopoietic cells except RBCs/platelets |
| **CD31 (PECAM-1)** | Adhesion molecule, angiogenesis | Endothelial cells, platelets, some leukocytes |
| **MMP9** | Matrix metalloproteinase, tissue remodeling | Neutrophils, macrophages, tumor cells |
| **CD20** | B-cell receptor component | Mature B cells |
| **CD204 (MSR1)** | Scavenger receptor, phagocytosis | Macrophages |
| **p53** | Tumor suppressor protein, apoptosis and DNA repair | All nucleated cells (mutation accumulation in tumors) |
| **CD3** | TCR complex component | Pan-T cells |
| **LAMP3** | Lysosome-associated protein, DC maturation marker | Dendritic cells |
| **CD11c** | Integrin αXβ2, adhesion, phagocytosis | Dendritic cells, some macrophages |

| | | |
|---|---|---|
| **PD1** | Immune checkpoint receptor | Exhausted/activated T cells, some B cells |
| **CD73** | Ecto-5'-nucleotidase, adenosine production, immunosuppression | Tregs, stromal cells, endothelial cells |
| **Bcl2** | Anti-apoptotic protein | Broadly expressed, survival in lymphocytes and tumor cells |
| **GATA3** | Transcription factor, Th2 differentiation | Th2 $CD4^+$ T cells, luminal epithelial cells |
| **CD155 (PVR)** | Adhesion/immune checkpoint ligand | Tumor cells, dendritic cells, fibroblasts |
| **CD10** | Endopeptidase, diagnostic marker | Germinal center B cells, neutrophils, epithelial cells |
| **NKG2A** | Inhibitory NK cell receptor | NK cells, some $CD8^+$ T cells |
| **FOXP3** | Transcription factor, regulatory program | Regulatory T cells (Tregs) |
| **CXCL13** | Chemokine attracting B cells | Follicular helper T cells, stromal cells |
| **CD8a** | Cytotoxic receptor chain | $CD8^+$ T cells |
| **EOMES** | Transcription factor, cytotoxic program | $CD8^+$ T cells, NK cells |
| **CD137 (4-1BB)** | Costimulatory receptor | Activated $CD8^+$ and $CD4^+$ T cells |
| **CD134 (OX40)** | Costimulatory receptor | $CD4^+$ T & $CD8^+$ T cells |
| **CD209 (DC-SIGN)** | C-type lectin receptor | Dendritic cells, macrophages |
| **CD56 (NCAM)** | Adhesion molecule, NK marker | NK cells, subset of T cells |
| **TBET (T-bet)** | Transcription factor, Th1 program | Th1 $CD4^+$ T cells, $CD8^+$ T cells, NK cells |
| **GITR** | Costimulatory receptor | Regulatory T cells, activated T cells |

| | | |
|---|---|---|
| **Ecad (E-cadherin)** | Adhesion molecule | Tumor / Epithelial cells |
| **Tim3** | Inhibitory checkpoint receptor | Exhausted/activated T cells, some myeloid cells |
| **CXCL8 (IL-8)** | Chemokine recruiting neutrophils | Tumor cells, macrophages, epithelial cells |
| **CD66b** | Granulocyte activation marker | Neutrophils |
| **DNA1** | DNA intercalator (nuclear segmentation) | All nucleated cells |
| **DNA2** | DNA intercalator (nuclear segmentation) | All nucleated cells |
| **Ki67** | Nuclear proliferation marker | Cycling cells |
| **Podoplanin (D2-40)** | Lymphatic endothelial marker, adhesion | Lymphatic endothelial cells, stromal fibroblasts |
| **IgG** | Immunoglobulin G, antibody isotype | Plasma cells, B cells (secreted form) |
| **CD15** | Adhesion molecule, granulocyte marker | Neutrophils, eosinophils |

**Extended Table 2: Panel 1 IMC Markers: Function and Cell Type**

| | All | Panel 1 | Panel 2 |
|---|---|---|---|
| **n** | **192** | **191** | **189** |
| **Age** (years) **Median** (0.25, 0.75) quantile | **66** (60, 72.5) | **66** (60, 72.5) | **66** (60, 73) |
| **Female** (%) | **79** (41%) | **79** (41%) | **79** (42%) |
| **TNM** (%) | | | |
| I | **72** (38%) | **72** (38%) | **72** (38%) |
| II | **55** (29%) | **55**(29%) | **52** (28%) |
| III | **56** (29%) | **55** (29%) | **56** (30%) |
| IV | **9** (5%) | **9** (5%) | **9** (5%) |
| **Histology** (%) | | | |
| Adenocarcinoma | **124** (65%) | **124** (65%) | **123** (65%) |
| Squamous cell carcinoma | **41** (21%) | **41** (21%) | **40** (21%) |
| Adenosquamous carcinoma | **6** (3%) | **6** (3%) | **6** (3%) |
| Other | **20** (10%) | **20** (10%) | **20** (11%) |
| **CPS Positive** (%) | **95** (49%) | **95** (50%) | **95** (50%) |
| **TPS Positive** (%) | **57** (30%) | **57** (30%) | **57** (30%) |
| **TMB** (%) | | | |
| < 10 | **89** (46%) | **89** (47%) | **89** (47%) |
| [10, 16) | **81** (42%) | **81** (42%) | **81** (42%) |
| >= 16 | **20** (10%) | **20** (10%) | **20** (11%) |
| **Dead** (%) | **42** (22%) | **42** (22%) | **42** (22%) |
| **OS Median days** (0.25, 0.75) quantile | **1023** (638.5, 1435) | **1023** (638.5, 1435) | **1024** (641, 1449) |
| **Smoking status** (%) | | | |
| Ex-smoker | **69** (36%) | **69** (36%) | **68** (36%) |
| Non-smoker | **65** (34%) | **65** (34%) | **64** (34%) |
| Current smoker | **57** (30%) | **57** (30%) | **57** (30%) |

**Extended Table 3: Cohort summary for Immucan NSCLC across IMC Panels.**

| | Histology | | | |
|---|---|---|---|---|
| | Adenocarcinoma (n=125) | Squamous Cell carcinoma (n=41) | Other (n=26) | Total (n=192) |
| | N (%) | N (%) | N (%) | N (%) |
| **Sex** | | | | |
| **Female** | 62 (50) | 9 (22) | 9 (34.6) | 80 (41.7) |
| **Male** | 63(50) | 32 (78) | 17 (65.4) | 112 (58.3) |
| **Age at diagnosis** | | | | |
| **Median** | 67 | 66 | 66 | 66 |
| **Range** | 35 - 85 | 42 - 84 | 18 - 78 | 18 - 85 |
| **Stage at diagnosis** | | | | |
| **Stage I** | 56 (44.8) | 6 (14.6) | 9 (34.7) | 71 (37) |
| **Stage II** | 29 (23.2) | 20 (48.8) | 7 (26.9) | 56 (29.2) |
| **Stage III** | 35 (28) | 14 (34.1) | 7 (26.9) | 56 (29.2) |
| **Stage IV** | 5 (4) | 1 (2.5) | 3 (11.5) | 9 (4.6) |
| **Treatment received** | | | | |
| **Neo-adjuvant** | 11 (8.8) | 1 (2.4) | 2 (7.7) | 14 (7.3) |
| **Curative surgery** | 125 (100) | 41 (100) | 25 (96.2) | 191 (99.5) |
| **Adjuvant** | 33 (26.4) | 25 (61) | 15 (57.7) | 73 (38) |
| **Metastatic line** | 26 (20.8) | 4 (9.8) | 9 (34.6) | 39 (20.3) |
| **Smoking status at diagnosis** | | | | |
| **Current** | 56 (44.8) | 23 (56.1) | 14 (53.8) | 93 (48.4) |
| **Former** | 18 (14.4) | 13 (31.7) | 3 (11.6) | 34 (17.7) |
| **Never** | 26 (20.8) | 0 (0) | 5 (19.2) | 31 (16.2) |
| **Unknown** | 25 (20) | 5 (12.2) | 4 (15.4) | 34 (17.7) |
| **Status** | | | | |
| **Alive / Lost to FU** | 103 (82.4) | 29 (70.7) | 18 (69.2) | 150 (78.1) |
| **Dead** | 22 (17.6) | 12 (29.3) | 8 (30.8) | 42 (21.9) |

**Extended Table 4: Cohort summary for Immucan NSCLC divided by histology.**

| CN | Min | **Median** (0.25, 0.75) quantile | Max | **Mean** (Std) | Total | Slides with count > **0**/10/100 | Patients with count > **0**/10/100 |
|---|---|---|---|---|---|---|---|
| **0** | **0** | **114** (67, 181) | **712** | **134.13** (93.19) | **76318** | **568**/560/320 | **191**/191/183 |
| **1** | **2** | **338** (244, 473) | **1245** | **374.64** (185.86) | **213171** | **569**/568/558 | **191**/191/191 |
| **2** | **0** | **490** (393, 617) | **2289** | **527.14** (209.82) | **299944** | **568**/568/568 | **191**/191/191 |
| **3** | **0** | **547** (450, 649) | **1345** | **565.08** (177.65) | **321528** | **568**/568/568 | **191**/191/191 |
| **4** | **0** | **491** (368, 641) | **1610** | **515.24** (206.28) | **293173** | **568**/568/566 | **191**/191/191 |
| **5** | **0** | **312** (198, 492) | **1839** | **366.87** (234.68) | **208747** | **568**/568/537 | **191**/191/191 |
| **6** | **0** | **105** (58, 183) | **1236** | **144.92** (135.95( | **82457** | **567**/564/293 | **191**/191/182 |

**Extended Table 5: Count statistics for Immucan NSCLC Panel 1 PLDA.1 Tumor CNs.**

| CN | Min | **Median** (0.25, 0.75) quantile | Max | **Mean** (Std) | Total | Slides with count > **0**/10/100 | Patients with count > **0**/10/100 |
|---|---|---|---|---|---|---|---|
| **0** | **0** | **145** (71, 244,5) | **831** | **165,67** (121,97) | **28496** | **171**/166/113 | **171**/166/113 |
| **1** | **0** | **80** (50,25, 138) | **374** | **102,10** (71,87) | **17562** | **171**/165/73 | **171**/165/73 |
| **2** | **0** | **245** (97,75, 422,75) | **1143** | **296,57** (250,74) | **51010** | **156**/154/127 | **156**/154/127 |
| **3** | **0** | **411,5** (224, 643,5) | **1982** | **499,51** (381,42) | **85915** | **171**/171/159 | **171**/171/159 |
| **4** | **0** | **20** (2, 98,75) | **1016** | **79,51** (150,17) | **13675** | **134**/98/43 | **134**/98/43 |
| **5** | **0** | **114,5** (26,75, 255,5) | **1466** | **181,13** (212,32) | **31155** | **163**/142/87 | **163**/142/87 |
| **6** | **0** | **0** (0, 6,5) | **564** | **21,16** (67,44) | **3639** | **75**/35/12 | **75**/35/12 |
| **7** | **0** | **77** (28, 195,5) | **736** | **131,03** (143,62) | **22537** | **164**/148/76 | **164**/148/76 |
| **8** | **0** | **6** (0, 45) | **702** | **48,73** (104,22) | **8381** | **120**/76/26 | **120**/76/26 |
| **9** | **6** | **75** (52, 107) | **202** | **82,06** (40,51) | **14115** | **172**/170/49 | **172**/170/49 |
| **10** | **0** | **0** (0, 0) | **549** | **15,33** (66,16) | **2637** | **27**/16/9 | **27**/16/9 |
| **11** | **0** | **145,5** (15,75, 322,25) | **1333** | **225,03** (258,73) | **38706** | **142**/134/100 | **142**/134/100 |
| **12** | **0** | **30,5** (7, 91) | **1116** | **88,65** (148,21) | **15247** | **152**/127/40 | **152**/127/40 |
| **13** | **0** | **8** (1, 40,75) | **1545** | **50,15** (141,21) | **8626** | **139**/78/24 | **139**/78/24 |
| **14** | **0** | **33,5** (14, 60,25) | **326** | **48,44** (52,42) | **8331** | **167**/141/22 | **167**/141/22 |
| **15** | **0** | **111** (21, 278,5) | **788** | **172,61** (179,93) | **29689** | **166**/144/89 | **166**/144/89 |
| **16** | **0** | **181,5** (122, 268) | **480** | **199,56** (107,69) | **34324** | **170**/170/139 | **170**/170/139 |
| **17** | **0** | **1** (0, 39,75) | **1011** | **64,38** (151,70) | **11074** | **91**/59/30 | **91**/59/30 |
| **18** | **0** | **2** (0, 37) | **705** | **43,85** (100,25) | **7542** | **96**/62/20 | **96**/62/20 |
| **19** | **0** | **239,5** (143,25, 381,5) | **1189** | **295,59** (222,51) | **50841** | **171**/169/143 | **171**/169/143 |
| **20** | **0** | **186** (67,75, 363,75) | **1428** | **280,66** (302,14) | **48273** | **168**/158/114 | **168**/158/114 |
| **21** | **0** | **301** (113, 474,25) | **1317** | **337,78** (270,01) | **58098** | **170**/166/136 | **170**/166/136 |
| **22** | **0** | **0** (0, 5,25) | **849** | **18,48** (76,26) | **3179** | **65**/35/7 | **65**/35/7 |

**Extended Table 6: Count statistics for Immucan NSCLC Panel 1 PLDA.75 CD20$^+$ CNs.**

| CN | Min | **Median** (0.25, 0.75) quantile | Max | **Mean** (Std) | Total | Slides with count > **0**/10/100 | Patients with count > **0**/10/100 |
|---|---|---|---|---|---|---|---|
| **0** | **0** | **179** (84, 264) | **692** | **188,06** (131,91) | **107006** | **558**/538/401 | **190**/189/180 |
| **1** | **0** | **135** (73, 219) | **750** | **161,25** (119,18) | **91753** | **567**/559/370 | **191**/191/180 |
| **2** | **0** | **0** (0, 0) | **121** | **1,43** (10,21) | **816** | **23**/12/2 | **20**/11/3 |
| **3** | **0** | **0** (0, 0) | **1539** | **12,87** (81,22) | **7321** | **104**/70/16 | **80**/62/15 |
| **4** | **0** | **32** (2, 179) | **1595** | **152,23** (261,59) | **86619** | **461**/352/188 | **183**/158/108 |
| **5** | **0** | **29** (2, 133) | **1941** | **119,69** (229,12) | **68106** | **460**/342/170 | **183**/159/108 |
| **6** | **0** | **6** (0, 58) | **2005** | **85,39** (200,89) | **48589** | **372**/248/113 | **159**/125/71 |
| **7** | **0** | **115** (27, 291) | **1484** | **191,91** (222,99) | **109194** | **534**/476/303 | **189**/183/152 |
| **8** | **0** | **31** (2, 139) | **2187** | **123,85** (225,21) | **70473** | **454**/355/185 | **180**/160/115 |
| **9** | **0** | **95** (64, 128) | **492** | **103,28** (57,62) | **58765** | **568**/566/250 | **191**/191/187 |
| **10** | **0** | **0** (0, 9) | **2221** | **85,33** (267,60) | **48550** | **183**/138/87 | **96**/77/52 |
| **11** | **0** | **0** (0, 0) | **281** | **0,95** (13,00) | **539** | **7**/6/1 | **6**/5/2 |
| **12** | **0** | **51** (8, 181) | **2946** | **140,03** (257,96) | **79679** | **509**/410/203 | **189**/180/142 |
| **13** | **0** | **38** (4, 121) | **1258** | **96,67** (144,07) | **53298** | **489**/376/170 | **185**/163/115 |
| **14** | **0** | **72** (30, 132) | **716** | **98,13** (94,28) | **55834** | **564**/515/208 | **191**/188/147 |
| **15** | **0** | **26** (3, 126) | **1159** | **88,52** (140,22) | **50368** | **476**/357/166 | **187**/161/109 |
| **16** | **0** | **185** (103, 285) | **751** | **205,99** (132,77) | **117211** | **565**/555434 | **191**/191/189 |
| **17** | **0** | **14** (0, 166) | **1616** | **136,20** (254,16) | **77499** | **395**/298/173 | **165**/136/93 |
| **18** | **0** | **25** (0, 146) | **1870** | **129,41** (139,06) | **73636** | **414**/330/177 | **168**/149/101 |
| **19** | **0** | **0** (0, 0) | **1082** | **10,63** (60,01) | **6048** | **132**/71/12 | **94**/63/8 |
| **20** | **0** | **68** (13, 230) | **1530** | **163,77** (224,59) | **93187** | **516**/435/240 | **189**/181/136 |
| **21** | **0** | **167** (54, 363) | **1362** | **246,95** (245,72) | **140513** | **550**/515/365 | **190**/189/173 |
| **22** | **0** | **3** (0, 42) | **3250** | **88,46** (254,86) | **50334** | **340**/215/95 | **149**/108/61 |

**Extended Table 7: Count statistics for Immucan NSCLC Panel 1 PLDA.75 Tumor CNs.**

| CN | Min | **Median** (0.25, 0.75) quantile | Max | **Mean** (Std) | Total | Slides with count > **0**/10/100 | Patients with count > **0**/10/100 |
|---|---|---|---|---|---|---|---|
| **0** | **0** | **0** (0, 0) | **2904** | **24.54** (234.05) | **13889** | **16**/13/7 | **8**/7/4 |
| **1** | **0** | **101** (41.25, 185) | **590** | **128.35** (111.84) | **72647** | **555**/516/284 | **188**/185/168 |
| **2** | **0** | **86** (14, 277.75) | **1458** | **193.18** (253.88) | **109339** | **515**/441/271 | **183**/172/142 |
| **3** | **0** | **6** (0, 30) | **3156** | **52.79** (200.26) | **29878** | **409**/236/58 | **174**/135/53 |
| **4** | **0** | **11.5** (0, 134.75) | **1914** | **129.70** (258) | **73408** | **378**/285/163 | **157**/136/99 |
| **5** | **0** | **0** (0, 0) | **2878** | **43.46** (260.26) | **24596** | **69**/45/26 | **37**/20/13 |
| **6** | **0** | **76.5**, (14, 186) | **590** | **112.79** (114.91) | **63840** | **511**/442/240 | **183**/171/140 |
| **7** | **0** | **0** (0, 2) | **1870** | **62.54** (210.85) | **35397** | **162**/108/68 | **79**/61/40 |
| **8** | **0** | **40.5** (4, 158.25) | **2198** | **142.76** (258.04) | **80800** | **477**/371/182 | **181**/166/131 |
| **9** | **0** | **0** (0, 39.75) | **2210** | **76.64** (218.89) | **43376** | **248**/193/96 | **135**/115/76 |
| **10** | **0** | **0** (0, 0) | **3594** | **61.03** (290.08) | **34545** | **110**/72/39 | **64**/41/26 |
| **11** | **0** | **0** (0, 0) | **4282** | **19.84** (267.44) | **11227** | **6**/6/5 | **2**/2/2 |
| **12** | **0** | **85** (18, 220) | **1422** | **156.15** (191.72) | **88379** | **531**/459/264 | **187/184/**153 |
| **13** | **0** | **5** (0, 67.75) | **2099** | **103.45** (242.01) | **58551** | **371**/245/124 | **167**/131/75 |
| **14** | **0** | **8** (0, 50.75) | **748** | **46.91** (93.70) | **26552** | **387**/261/79 | **162**/129/69 |
| **15** | **0** | **100** (24.25, 213) | **1285** | **146.24** (168.34) | **82771** | **529**/464/280 | **186**/177/147 |
| **16** | **0** | **2.5** (0, 23) | **2062** | **59.84** (205.80) | **33872** | **350**/185/63 | **161**/109/45 |
| **17** | **0** | **1** (0, 41.5) | **2261** | **85.48** (258.10) | **48381** | **305**/212/93 | **150**/116/71 |
| **18** | **0** | **78.5** (20, 183) | **2009** | **141.83** (198.29) | **80274** | **530**/460/249 | **185**/174/151 |
| **19** | **0** | **0** (0, 0) | **1638** | **24.35** (134.98) | **13780** | **73**/48/26 | **41**/27/15 |
| **20** | **0** | **0** (0, 1) | **1689** | **30.09** (145.82) | **17032** | **144**/89/34 | **83**/59/27 |
| **21** | **0** | **98** (18, 264.75) | **1142** | **170.91** (199.63) | **96736** | **518**/454/275 | **183**/171/153 |
| **22** | **0** | **0** (0, 7) | **1908** | **63.40** (197.41) | **35886** | **164**/137/90 | **107**/99/72 |
| **23** | **0** | **0** (0, 0) | **3125** | **53.22** (288.81) | **30122** | **72**/53/35 | **34**/28/19 |
| **24** | **0** | **0** (0, 0) | **2142** | **26.42** (156.56) | **14955** | **89**/54/26 | **61**/36/15 |
| **25** | **0** | **20** (10, 41.75) | **200** | **30.95** (30.90) | **17516** | **549**/414/23 | **188**/179/63 |
| **26** | **0** | **136** (51, 292) | **1262** | **196.09** (199.25) | **110988** | **540**/498/340 | **188**/184/169 |
| **27** | **0** | **88.5** (17, 242.5) | **1002** | **156.69** (179.44) | **88687** | **521**/451/267 | **186**/180/149 |
| **28** | **0** | **0** (0, 0) | **2211** | **34.42** (200.86) | **19481** | **97**/51/27 | **54**/29/16 |
| **29** | **0** | **0** (0, 0) | **2515** | **22.12** (176.78) | **12521** | **26**/19/10 | **16**/11/5 |
| **30** | **0** | **12** (1, 65) | **1266** | **64.76** (140.70) | **36656** | **437**/291/100 | **183**/152/82 |
| **31** | **0** | **4** (0, 87.75) | **3033** | **126.59** (302.74) | **71650** | **355**/233/138 | **158**/125/87 |
| **32** | **0** | **0** (0, 0) | **5169** | **63.45** (464.70) | **35910** | **23**/20/16 | **10**/9/8 |
| **33** | **0** | **1** (0, 33.75) | **3006** | **75.95** (258.72) | **42988** | **320**/203/90 | **147**/113/66 |

**Extended table 8: Count statistics for Immucan NSCLC Panel 2 PLDA.100 CNs.**

| CN | Min | **Median** (0.25, 0.75) quantile | Max | **Mean** (Std) | Total | Slides with count > **0**/10/100 | Patients with count > **0**/10/100 |
|---|---|---|---|---|---|---|---|
| **0** | **0** | **21** (4, 58) | **929** | **46.71** (82.06) | **17237** | **309**/238/53 | **309**/238/53 |
| **1** | **0** | **24** (5, 74) | **1027** | **71.95** (132.32) | **26549** | **306**/245/69 | **306**/245/69 |
| **2** | **0** | **0** (0, 11) | **2245** | **65.23** (224.83) | **24071** | **150**/94/42 | **150**/94/42 |
| **3** | **0** | **28** (8, 93) | **920** | **76.68** (121.16) | **28295** | **325**/254/83 | **325**/254/83 |
| **4** | **0** | **48** (17, 101) | **935** | **76.73** (95.55) | **28312** | **361**/312/95 | **361**/312/95 |
| **5** | **0** | **9** (0, 91) | **2022** | **105.25** (241.57) | **38838** | **245**/178/86 | **245**/178/86 |
| **6** | **0** | **111** (25, 251) | **1138** | **173.38** (189.99) | **63979** | **333**/300/196 | **333**/300/196 |
| **7** | **0** | **66** (44, 104) | **1345** | **92.27** (111.85) | **34049** | **369**/367/95 | **369**/367/95 |
| **8** | **0** | **0** (0, 0) | **3592** | **46.38** (268.03) | **17116** | **67**/36/21 | **67**/36/21 |
| **9** | **0** | **9** (0, 57) | **3359** | **93.62** (312.7) | **34545** | **237**/177/68 | **237**/177/68 |
| **10** | **0** | **75** (14, 224) | **910** | **151.66** (189.04) | **55961** | **326**/289/157 | **326**/289/157 |
| **11** | **0** | **64** (22, 156) | **1132** | **116.14** (154.22) | **42856** | **350**/313/129 | **350**/313/129 |
| **12** | **0** | **0** (0, 41) | **3089** | **98.98** (315.43) | **36525** | **167**/128/67 | **167**/128/67 |
| **13** | **0** | **2** (0, 28) | **3303** | **74.96** (279.75) | **27662** | **196**/123/50 | **196**/123/50 |
| **14** | **0** | **93** (38, 203) | **1069** | **140.85** (147.26) | **51974** | **366**/342/174 | **366**/342/174 |
| **15** | **0** | **0** (0, 17) | **2412** | **56.66** (201.32) | **20908** | **152**/105/40 | **152**/105/40 |
| **16** | **0** | **40** (9, 104) | **1258** | **88.58** (144.07) | **32685** | **322**/268/96 | **322**/268/96 |
| **17** | **0** | **73** (31, 134) | **768** | **100.29** (100.74) | **37006** | **357**/331/134 | **357**/331/134 |
| **18** | **0** | **2** (0, 24) | **1972** | **53.4** (175.25) | **19706** | **187**/133/41 | **187**/133/41 |
| **19** | **0** | **34** (7, 98) | **725** | **67.72** (93.45) | **24988** | **313**/263/91 | **313**/263/91 |
| **20** | **0** | **6** (0, 44) | **2527** | **67.3** (212.88) | **24835** | **234**/163/58 | **234**/163/58 |
| **21** | **0** | **23** (5, 85) | **2766** | **87.8** (216.56) | **32397** | **318**/238/83 | **318**/238/83 |
| **22** | **0** | **54** (18, 134) | **975** | **106.98** (135.27) | **39476** | **354**/313/126 | **354**/313/126 |
| **23** | **0** | **0** (0, 36) | **1285** | **63.89** (173.78) | **23577** | **156**/119/52 | **156**/119/52 |
| **24** | **0** | **69** (25, 148) | **746** | **105.73** (116.33) | **39014** | **331**/309/141 | **331**/309/141 |
| **25** | **0** | **25** (0, 76) | **1025** | **57.64** (94.27) | **21268** | **273**/219/68 | **273**/219/68 |
| **26** | **0** | **0** (0, 0) | **3664** | **32.08** (253.18) | **11839** | **64**/33/15 | **64**/33/15 |

**Extended table 9: Count statistics for Jackson et al. IMC PLDA.100 CNs.**
